# Introgression under linear selection on continuous genomes

**DOI:** 10.64898/2026.01.30.702779

**Authors:** Félix Foutel-Rodier, Nick Barton, Alison Etheridge

## Abstract

We model the introgression of a small block that carries many weakly selected linked loci into a large homogeneous population, under the simple assumption that a block of introduced genome has a selective effect proportional to its map length. Using a diffusion approximation, we compute the probability that some part of the initial block survives the initial phase of the introgression and give the typical length of the surviving blocks. Our results quantify the effect of recombination on selection and drift during an introgression and indicate that the fate of the block depends on the strength of selection relative to recombination. When selection is positive some parts of the block are able to survive at large times, but large blocks can only persist if selection is stronger than recombination. Surprisingly, the probability of such a successful introgression is independent of the strength of recombination and is the same as that for a single beneficial allele. (This is not true of other quantities.) Conversely, a deleterious or neutral block is eventually lost, but at a much slower rate than a single allele with the same selective effect. In this case, surviving blocks are very small. We also consider the introgression of a block of genome made of a single beneficial allele linked to a deleterious background and compute the amount of deleterious material that hitchhikes during fixation.

## 1 Introduction

Inherited variation in quantitative traits is typically due to very many genetic loci. This is implied by the sustained response to artificial selection, which typically shifts traits by many standard deviations, with change continuing over a hundred or more generations. Over the past decade, the highly polygenic basis of heritable variation has been confirmed by large genome-wide association studies, which routinely identify an extraordinary number of variants, spread across the genome. Indeed, practical quantitative genetics is based on the infinitesimal model, which assumes an indefinite number of unlinked loci (Barton et al., 2017). Of course, this multitude of variants must be spread across a genome of finite map length. Thus, at least in the short term, selection must act on the aggregate effects of blocks of genome, rather than on individual variants. Very many generations of recombination are needed to disentangle these variants, especially if there are multiple causal alleles, tightly linked within functional genes.

Compared with the elaborate population genetics of discrete loci, the theory for linked blocks of genome is relatively undeveloped. Fisher (1954) introduced the theory of junctions, to describe the inbreeding of continuous genomes, whilst Slatkin (1972) and Robertson (1977) analysed selection on whole chromosomes. Somewhat more recently, there has been substantial work on ‘background selection,’ where the elimination of deleterious mutations amplifies random drift (Charlesworth et al., 1993); Hudson and Kaplan (1995) gives an expression for an arbitrary number of loci with multiplicative effects, which shows that background selection acts primarily through the rate of mutation per map length, *U/R*. Good et al. (2014) identified a regime where the key variable is the fitness variance per map length. Such effects of linked selection can select for modifiers of recombination. Barton (1995) gave general expressions for the selection on recombination in an infinite population, in terms of the mean and variance of log fitness; Roze (2021) extended these to the case of a finite population, considering Hill–Robertson interference due to deleterious mutations. These theoretical results are mostly derived using theory and simulations for discrete loci, but the results depend on the aggregate effects of selection across the genome, suggesting that they could be derived from an infinitesimal limit.

Here, we consider a simple case: the fate of a single block of genome, introduced into a large (but finite) homogeneous population. The case of an infinite population is straightforward, and also covers the expected contribution of a single block. Specifically, introgression is reduced by selection against deleterious linked variants (or increased if selection favours them). With no linkage, the reduction is just the product of mean fitnesses in successive backcross generations, whilst with linkage, the reduction in gene flow can be found explicitly if selection is additive (Westram et al., 2022, SI2, Eq 5), which can readily be extended to include epistasis. For a variety of models, the net effect depends primarily on the relative rates of selection versus recombination.

The stochastic case is more intricate, but is tractable in the early stages of introgression, when individuals carrying introduced material are rare, and therefore reproduce independently, in a branching process. Baird et al. (2003) consider the neutral case, and show that although an individual block of genome will eventually be entirely lost, this is a very slow process: if the block has map length *y*, the probability of fragments being present after *t* generations decays as ∼*y/* log(*ty/*2), in contrast to ∼1*/t* for a single neutral locus. Small fragments persist for a very long time and may become common. Baird et al. (2003) derive the generating function for this process, allowing calculation of the mean and variance of numbers and sizes of introgressing blocks.

Sachdeva and Barton (2018a) define the infinitesimal model with linkage: each genome carries an indefinitely large number of variants, so that any block has a net effect that is drawn from a Gaussian, with zero mean and variance proportional to its length; as in Robertson (1977), smaller blocks have effects also drawn from a Gaussian. Sachdeva and Barton (2018a) show that under directional selection, blocks that have a selective advantage greater than their map length can increase intact, throwing off smaller subblocks as they do so.

Here, we extend Baird et al. (2003) to the simplest case of linear selection, where a block of genome has an effect on fitness proportional to its length. This is considerably simpler than the infinitesimal model with linkage (Robertson, 1977; Sachdeva and Barton, 2018a), because the effect of a block is entirely determined by its length. Both the model considered here and that of Sachdeva and Barton (2018a) arise in the limit of a genome with a large number *L* of variants, each of small effect, as *L*→ ∞. Whereas in the model of Sachdeva and Barton (2018a) the mean effect of each variant is taken to be *o*(1*/L*) (i.e. it converges to zero faster than 1*/L* as *L* → ∞) and the variance of the effect is *O*(1*/L*), for the model we consider here the mean is *O*(1*/L*) and the variance is *o*(1*/L*). Thus, the relative magnitude of the mean and variance of effects tunes between the case of linear selection (studied here), and the infinitesimal with linkage, as defined by Sachdeva and Barton (2018a). Biologically, one can imagine a genome drawn from a highly diverged population, such that the effects of any block are uniformly deleterious or advantageous, in proportion to block length. Effects would be deleterious if the donor population were adapted to a different environment, or if it had accumulated many deleterious mutations. Conversely, effects would be positive if the recipient population had accumulated deleterious mutations, or if the two populations had different sets of (partially) recessive deleterious mutations, leading to heterosis.

Just as Baird et al. (2003) quantifies the ability of linkage to slow the eradication of (neutral) introgressing material through genetic drift, our results here will quantify the extent to which linkage counters the combined effects of selection and drift on an introgressing block. For example, we shall see that, even when the introduced block is uniformly deleterious, the probability that some introgressed material persists in the population after *t* generations declines polynomially in *t*, in contrast to the exponential decay for a single deleterious locus.

Following Baird et al. (2003), we introduce a branching process which encodes the length of the block of introgressed material carried by each individual (where an individual carrying no such material is deemed to have died). We then take a diffusion approximation, valid when selection and recombination are slow, and use its generating function to find simple expressions for the probability that some of the initial block will survive up to some given time; for the mean number and lengths of introgressing blocks; for the expected total amount of introgressed material as a function of time; and for the time to common ancestry of different blocks, which determines how closely they are clustered along the genome. To complement our calculation for the probability of survival of some introgressed material, no matter where on the genome, we also study the probability that there will be a block containing a specified locus at large times. Finally we consider the case in which a discrete beneficial allele is embedded in a block that has otherwise deleterious effects, and find how linked deleterious material retards its progress. These results depend primarily on the relative densities of selection relative to recombination; we discuss how one might estimate the strength of selection from the observed distributions of sizes of introgressing blocks.

## 2 Methods

### 2.1 The model

We consider a haploid population of fixed size *N* in which each individual expresses a one-dimensional trait, which is the sum of the contributions of *L* » 1 different genes uniformly distributed over a block of length *x*. Initially, the population is composed of genetically identical individuals (whose trait value is set to *z* = 0) and we introduce a single individual from a diverged population. The diverged population is assumed to be at linkage equilibrium, so that the contributions 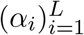 of the *L* loci to the trait are independent. We suppose that they are also identically distributed. We consider a regime in which Var(*α*_*i*_) « E[*α*_*i*_] and *L*E[*α*_*i*_] → *sx* ∈ ℝ as *L* → ∞. Under this assumption, the law of large numbers ensures that a block of length *y* has an effect *z*(*y*) = *sy* on the trait, as long as the number of loci that it carries is large, that is *yL* » 1. Note that this regime is different from that considered in Sachdeva and Barton (2018a), where it is assumed that |E[*α*_*i*_]| « Var(*α*_*i*_), *L* Var(*α*_*i*_) → *σ*^2^*x* and the fitness of blocks is described by a Brownian path.

We assume that the trait is under directional selection, and that the fitness of an individual with trait *z* is *e*^*z*^. We fix a parameter *r >* 0 with *rx <* 1. Generations are non-overlapping. In each generation, *N* pairs of parents are sampled proportional to fitness from the previous generation. For each such pair, independently, with probability 1 − *rx* there is no recombination within the block, and with probability *rx* a single crossover takes place at a point sampled uniformly in [0, *x*]. In either case this results in two possible haplotypes and the single offspring corresponding to this pair inherits each with equal probability. For small *rx* the assumption of a single crossover is a close approximation to more realistic maps, with a varying degree of interference between crossovers. For long blocks, one could simulate the process for a few generations, until introgressed blocks are short and well-separated on the genome, at which point they will evolve independently and our analysis applies to any one of those blocks.

### 2.2 Branching process approximation

We track the early dynamics of the genetic descendants of the introduced block, assuming that their number is negligible compared to the total population size *N*. The number of times that an individual carrying a block of introgressed material of length *y* is among the two parents chosen to produce an offspring in the next generation will be approximately distributed as a Poisson(2*e*^*sy*^) random variable. With high probability, in this branching phase, each time it is selected, the other parent in the pair does not carry introgressed material. If there is no recombination event, or if the crossover point lies outside the introgressed block (events with total probability 1 − *ry*), then with probability 1*/*2 the offspring inherits the whole block of length *y*, and with probability 1*/*2 it inherits none of the introgressed material. On the other hand, with probability *ry* a crossover event falls at a uniformly distributed point within the block, and so the offspring inherits a subblock of introgressed material whose length is distributed as *Uy*, where *U* ∼ Uniform(0, 1). As long as the number of individuals carrying some introgressed material is negligible, these steps can be assumed to occur independently for different individuals.

Collecting these steps, we are led to consider a branching process that records the length of the blocks of introgressed material. It starts from a single ancestor with block length *x*. An individual carrying a block of length *y* gives birth to

- a Poisson (*e*^*sy*^(1 − *ry*)) distributed number of offspring with block length *y*; and
- a Poisson (2*e*^*sy*^*ry)* distributed number of offspring with independent block lengths distributed as *Uy*, where *U* ∼ Uniform(0, 1).

### 2.3 Diffusion approximation

We shall assume that the parameters *r* and *s* of our model are small, and the proofs of our results will use a diffusion approximation to the branching process described above. This is entirely analogous to the Feller diffusion approximation to a near-critical branching process, which is routinely used for analysis of a single genetic locus. However, because we must record the amount of introgressed material carried by each individual, instead of a single number recording the population size, we must track the sizes of the subpopulations carrying each possible block length in (0, *x*]. The mathematics becomes somewhat technical and for the most part is deferred to Section A.1. Here we motivate the form of the equations, but this section is not necessary for understanding the statements of our results in Section 3.

Classically we approximate the dynamics of a branching process in which the number of offspring of each individual has mean 1 + *γ/K* + *o*(1*/K*) and variance *η*^2^ + *O*(1*/K*), over timescales of order *K* generations (where *K* is some large parameter), by a Feller diffusion. Measuring population size in units of *K*, as *K* → ∞ we find

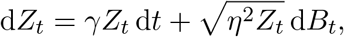

where *B*_*t*_ is Brownian motion. We can then approximate the original branching process at time *Kt* by *KZ*_*t*_ where *Z*_0_ = 1*/K*.

In our setting, the procedure above corresponds to taking *r* = *ρ/K* and *s* = *σ/K*. Consider the size of the population that carries blocks with lengths in the infinitesimal interval [*y, y*+d*y*) for *y* ∈ (0, *x*]. If an individual carries a block of length *y*, then the number of its offspring that inherit the entire block has mean *e*^*sy*^(1 − *ry*) = 1 +(*σ* − *ρ*)*y/K* +*o*(1*/K*) and variance 1 + *O*(1*/K*). This is consistent with the conditions required to obtain a Feller diffusion as *K* → ∞. Blocks with lengths falling in [*y, y* + *dy*) can also arise through recombination of a larger block. The number of offspring of an individual carrying a block of length *z > y* that carry a subblock of the parental block has mean 2*ρz/K* + *o*(1*/K*) and variance *O*(1*/K*). For each such offspring (independently) the chance that the subblock has length in the interval [*y, y* +d*y*) is d*y/z*, and so (the *z*’s cancel and) over time intervals of *O*(*K*) generations, we see an influx to the population carrying blocks in [*y, y* + d*y*) at rate 2*ρ* times the size of the population carrying blocks in [*z, z* + *dz*), for each *z* ∈ (*y, x*].

By computing the infinitesimal mean and variance of the branching process, we argue in Section A.1 that, if the population starts from *K* blocks with initial length *x*, measuring both population size and time in units of size *K*, as *K* → ∞ the size of the population at time *t* that carries a block of introgressed material with length in [*y, y* + d*y*) converges to the solution to the stochastic partial differential equation

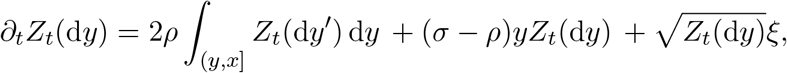

started from *Z*_0_(d*y*) = *δ*_*x*_(*y*) (an atom of mass one at *x*) and where *ξ* is a space-time white noise. The integral term corresponds to the formation of smaller blocks due to the erosion of longer blocks by recombination. The other two terms indicate that the number of copies of a block of length *y* fluctuates according to a Feller diffusion with mean *γ* = (*σ* − *ρ*)*y* and variance *η*^2^ = 1, and that those diffusions are independent for different block lengths. Formally, the solution to this equation is defined by considering the dynamics of the integral of *Z*_*t*_ against a test function, see Section A.1.

In the calculation above, the number of offspring of an individual was taken to be Poisson distributed. More generally, suppose that the number of ‘parental pairs’ in which an individual is involved is given by the random variable *P* with mean 2 + *O*(1*/K*) and variance *c*^2^ + *O*(1*/K*), and for each offspring, independently, whether it inherits some of the introgressed material is determined by a Bernoulli random variable *χ* which takes the value 1 with probability 1*/*2+ *O*(1*/K*) and zero otherwise. Then the variance of the total number of offspring carrying any of the block is E[*P*] Var(*χ*) +Var(*P*)E[*χ*]^2^ = (*c*^2^ + 2)*/*4 + (1*/K*). The parameter *η*^2^ = (*c*^2^ + 2)*/*4 then appears as a coefficient in the noise (corresponding to *N/N*_e_), exactly as in the Feller diffusion, and our equation becomes

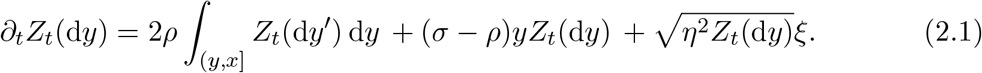

The theoretical expressions in Section 3 are obtained by using the solution of (2.1) to approximate the branching process. More precisely, the branching process started from a single block of length *x* is approximated at times *Kt*, for *K* large, by *KZ*_*t*_ where *Z*_0_ = *δ*_*x*_*/K*. As becomes evident in the proofs provided in the appendices, our results for the original branching process can all be expressed independently of *K*; see in particular the discussion at the beginning of Section A.2 for more detail of how this works. Nonetheless, by using the diffusion approximation we are implicitly assuming that *r* and |*s*| are small, and that we are considering the process at large times.

### 2.4 Beneficial allele in a deleterious background

We shall apply the same approach to a second situation, in which the block of genome being introduced carries one beneficial allele linked to deleterious material. The total length of the initial block will still be denoted by *x*, and we assume that the location of the beneficial allele on that block is such that the amount of deleterious material linked to its left (resp. right) is *x*_1_ (resp. *x*_2_), with *x*_1_ + *x*_2_ = *x*. A subblock of length *y* has an additive effect *s*_+_ + *ys*_−_ on the trait if it contains the beneficial allele, and *ys*_−_ otherwise, with *s*_+_ *>* 0, *s*_−_ *<* 0. All other aspects of the model are kept identical.

We only keep track of the blocks that contain the positive allele. Each such block is recorded as a pair (*y*_1_, *y*_2_) giving the length of deleterious material to the left and to the right of the positive allele, respectively. We can apply the same steps as in the first situation: take a branching approximation of the Wright–Fisher dynamics and then a diffusion approximation of the branching process. In order for the diffusion approximation to be valid, we make the same assumptions on *s*_−_ and *r* as before, and now we add the additional assumption that *s*_+_ = *σ*_+_*/K* for some *σ*_+_ *>* 0. Measuring population size and time in units of *K* generations, in much the same way as before, the population can be approximated by the solution to a stochastic partial differential equation. The form of the equation is intuitively clear: for each pair (*y*_1_, *y*_2_) we expect a term that resembles the Feller diffusion, but now there is an extra term *σ*_+_*Z*_*t*_ reflecting the presence of the beneficial allele in the block. Smaller blocks will be created from larger blocks by recombination, but now, since we are only retaining a block of introgressed material if it also contains the beneficial locus, the rate will be proportional to *ρ* instead of 2*ρ*. To formulate the limit precisely, we introduce the following notation: for a measure *µ*(d*y*_1_, d*y*_2_) on [0, *x*_1_] × [0, *x*_2_], we define *µ*((*y*_1_, *x*_1_], d*y*_2_) to be the measure on [0, *x*_2_] such that

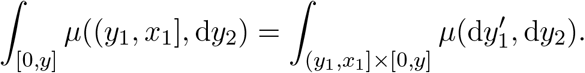

This will be used to represent the rate at which a block of size *y*_1_ to the left of the beneficial allele is ‘fed’ by recombination of longer blocks on the left. Similarly,

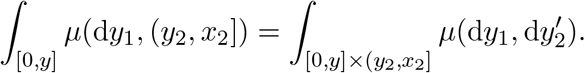

In this notation, the diffusion approximation is the solution to the stochastic partial differential equation

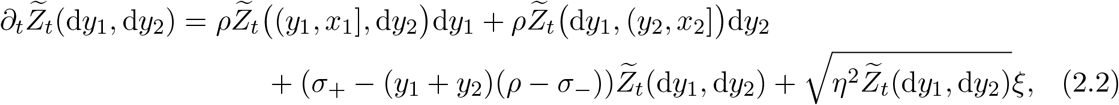

where *ξ* is a two-dimensional space-time white noise. This result can be derived using similar arguments to those outlined in Section A.1.

### 2.5 Numerical simulations

#### Simulation procedure

For *s < r*, the simulations were carried out as follows. The population in generation *t* is recorded as a collection of triples (*N*_*i,t*_, *L*_*i,t*_, *R*_*i,t*_)_*i*_, with the interpretation that there are *N*_*i,t*_ individuals carrying the same block corresponding to the interval [*L*_*i,t*_, *R*_*i,t*_]. At *t* = 0 we start from a single element (1, 0, *x*). Generation *t* + 1 is obtained as follows. For each *i*, we sample two independent random variables:

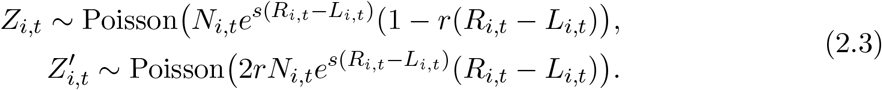

The first random variable corresponds to the number of descendants carrying exactly the block [*L*_*i,t*_, *R*_*i,t*_] and we add to the next generation (*Z*_*i,t*_, *L*_*i,t*_, *R*_*i,t*_). The second random variable corresponds to the number of descendants experiencing a crossover in their parent’s block. For each 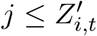, we let *U*_*i,t*_ be uniformly distributed over [*L*_*i,t*_, *R*_*i,t*_] and add to the population either (1, *U*_*i,t*_, *R*_*i,t*_) or (1, *L*_*i,t*_, *U*_*i,t*_) with equal probability.

This step is repeated until the population dies out, or *t* exceeds some specified final time, or the population size (defined as *N*_*t*_ = ∑_*i*_ *N*_*i,t*_) exceeds some specified threshold.

#### Culling procedure

When *s > r*, the population size grows faster than exponentially and we use a culling procedure to reduce the computational time. It depends on an additional parameter *N*_cul_. The first distinction from the approach for *s < r* is that we directly record the population in generation *t* as a collection of block lengths *Y*_*i,t*_, *i* = 1, …, *N*_*t*_ (with possible duplicates among the block lengths). The next generation is formed as follows. Independently for each *i*, let *Z*_*i,t*_ and 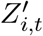 be two random variables as in (2.3), with *N*_*i,t*_ = 1 and *R*_*i,t*_ − *L*_*i,t*_ replaced by the block length *Y*_*i,t*_. Then, we add to the next generation *Z*_*i,t*_ blocks of length *Y*_*i,t*_, and 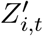 blocks whose lengths are independent and uniformly distributed over (0, *Y*_*i,t*_).

This step constructs a new collection of 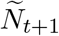 block lengths denoted by 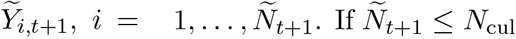, all these blocks are kept and form the next generation (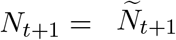 and 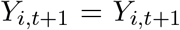). Otherwise, if 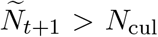, the next generation is obtained by sampling *N*_cul_ blocks from the vector 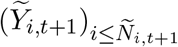, without replacement. (In particular in this case *N*_*t*+1_ = *N*_cul_.)

This step is repeated until the population dies out or *t* exceeds some specified final time. The process is not stopped when the population size hits the threshold *N*_cul_.

#### Data availability statement

Python codes used to generate the simulated data can be found at https://src.koda.cnrs.fr/3652jw85/linear-selection-introgression. The authors state that all data necessary for confirming the conclusions presented in the article are represented fully within the article.

## 3 Results

### 3.1 Examples of introgression

We start by discussing the behaviour of the branching process approximation for some examples. Figure 1 shows four realisations of the branching process (with no major effect locus), one for each of four values of the selection parameter (*s* = −0.01; 0; 0.01; 0.04), with *r* = 0.02. All simulations are conditioned on surviving until *t* = 5,000 generations or are stopped when the population reaches 50,000 blocks. Each line of the plot represents an individual, and the subblock that it carries is represented by the shaded region. The *y*-axis is the index of an individual and ranges from 1 to the total number of blocks; the *x*-axis represents the position along the initial block (which has length *x* = 1). Some additional independent realisations can be found in Figures S1 to S4.

**Figure 1:**
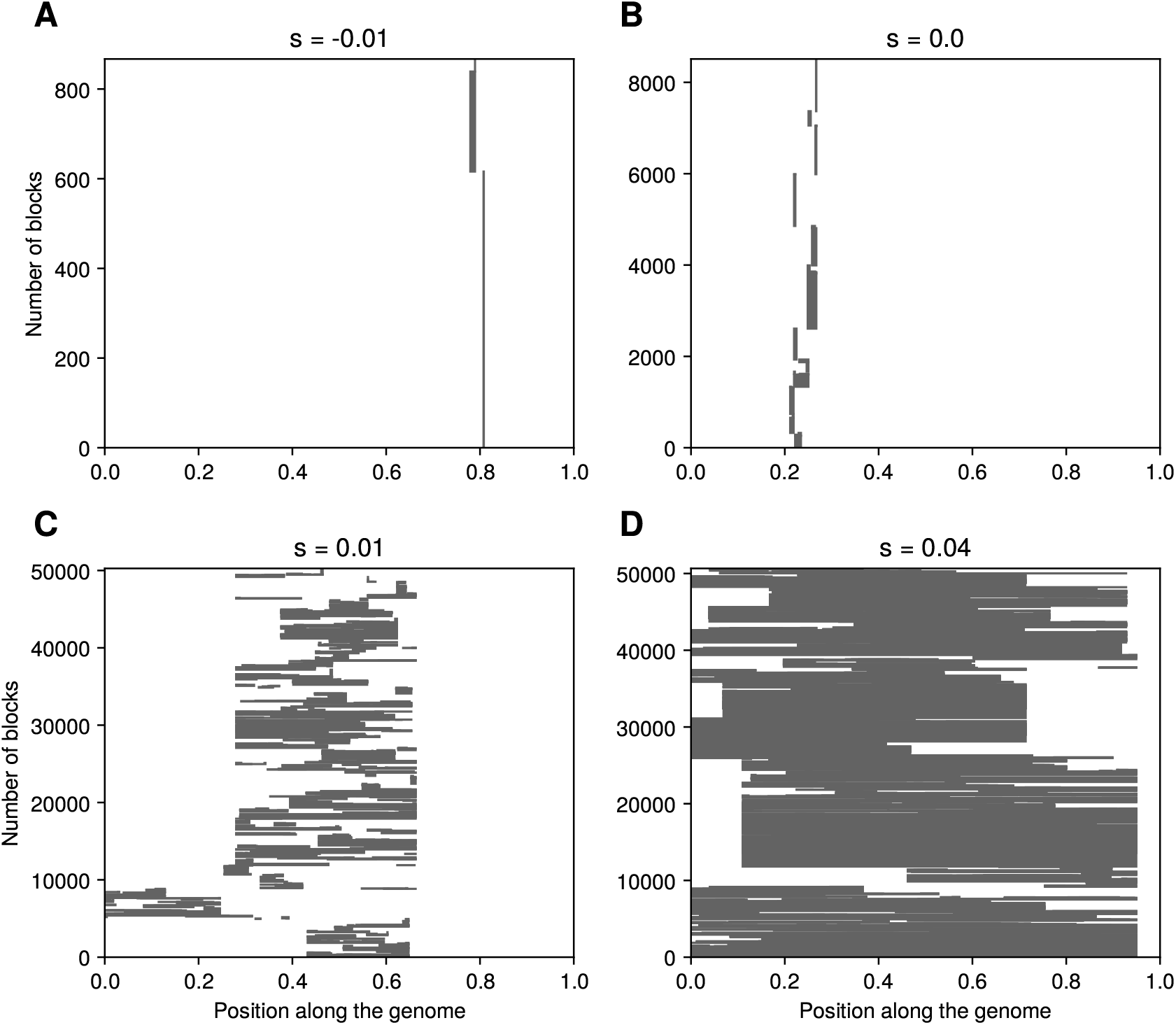
Realisations of the branching approximation. The process is stopped at generation *t* = 5,000 or when it reaches 50,000 blocks. In all simulations, *r* = 0.02 and the length of the initial block is *x* = 1. The process reached 50,000 blocks only for *s* = 0.01 and *s* = 0.04, after 1, 304 and 263 generations respectively.

We observe that the typical length of the blocks increases with the selective parameter *s*. For neutral (*s* = 0) and negative (*s <* 0) selection, the surviving blocks are very small and all concentrated in a narrow genomic region. The number of blocks does not reach 50,000 before generation *t* = 5,000, indicating that (conditional on establishment) the population size grows slowly. The population for *s* = −0.01 is made of copies of very few different blocks, whereas that for *s* = 0 is more diverse. When selection is positive (*s >* 0) blocks are found across a wide genomic region that almost spans the entire initial block. We observe that large blocks can survive when *s* = 0.04, whereas most surviving blocks are small when *s* = 0.01, with a few larger blocks (recall that *r* = 0.02). In both cases the population reaches 50,000 blocks rapidly, indicating that the number of blocks grows faster than for neutral and negative selection.

Our aim now is to provide a theoretical understanding of these observations by computing several quantities of interest (survival probability, mean number of blocks, and mean block length distribution). Since, as advertised, we are going to use the diffusion approximation, our results require that |*s*| « 1, *r* « 1, *rt* » 1, and that *r* is not negligible with respect to *s*. If *r* = 0, then the block evolves as a single locus with fitness effect *sx*, for which the theory is well established. Our results are summarised in Table 1. The results in Sections 3.2 to 3.7 deal with the model described in Section 2.1, with no major allele. The results for the model of Section 2.4 are given in Section 3.8.

**Table 1.** Summary of the principal theoretical results, with *p*_*t*_(*x*) the survival probability, *M*_*t*_(*x*) the expected number of blocks, *m*_*t*_(*x, y*) the expected number of blocks of length *y*, and *β* = (*r* + *s*)*/*(*r* − *s*). All the results hold for *rt* » 1 and *r, s* « 1.

| Regime | $\beta$ | $p_t(x)$ | $M_t(x)$ | $\int_y^x m_t(x, z)dz/M_t(x)$ |
| --- | --- | --- | --- | --- |
| $s > r$ | $(-\infty, -1)$ | $\frac{2sx}{\eta^2} - \frac{2(sx)^{\frac{1}{\beta}}}{\eta^2 t^{1-\frac{1}{\beta}}}$ | $\frac{((s-r)xt)^{-\beta-1}}{\Gamma(-\beta)} e^{(s-r)xt}$ | $\left(1 - \frac{y}{x}\right)^{-\beta-1}$ |
| $r > s > 0$ | $(1, \infty)$ | $\frac{2sx}{\eta^2} - \frac{2(sx)^{\frac{1}{\beta}}}{\eta^2 t^{1-\frac{1}{\beta}}}$ | $\frac{((r-s)xt)^\beta}{\Gamma(1+\beta)}$ | $e^{-(r-s)yt}$ |
| $s = 0$ | 1 | $\frac{rx}{\eta^2 \log(\frac{rxt}{2})}$ | $rx t$ | $e^{-(r-s)yt}$ |
| $s < 0$ | $(-1, 1)$ | $\frac{2(-sx)^\beta}{\eta^2 t^{1-\beta}}$ | $\frac{((r-s)xt)^\beta}{\Gamma(1+\beta)}$ | $e^{-(r-s)yt}$ |

### 3.2 Probability of introgression

Our first results concern the probability that some part of the initial block survives at large times. Let *p*_*t*_(*x*) be the probability that an introduced block of initial length *x* has genetic descendants alive in generation *t*. In Section A.2 we compute the limit of *p*_*t*_(*x*) by studying the generating function of the diffusion approximation. We find that the probability of survival depends on whether the introduced material has a selective effect

which is negative, neutral, or positive through the following characteristic exponent

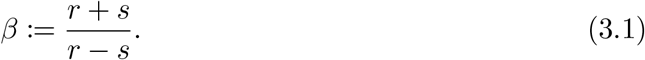

Note that *β* depends only on the relative rates of selection and recombination. It is sketched in Figure 2.

**Figure 2.**
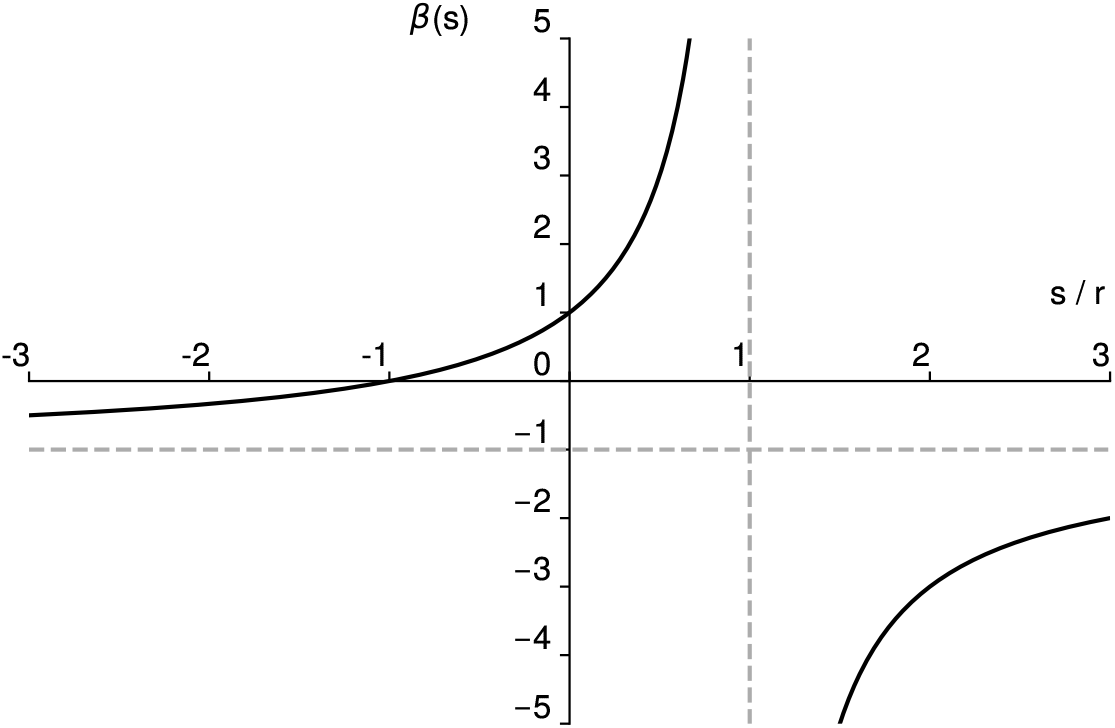
Sketch of *β* in (3.1) as a function of *s/r*.

#### Negative selection (*s <* 0)

When the selective effect is negative (*s <* 0 and so −1 *< β <* 1), we derive in Section A.2 that the survival probability decays as

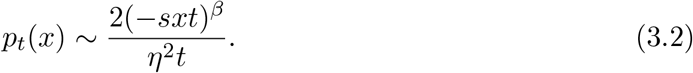

We remind the reader that in using asymptotics of the diffusion approximation we are assuming that *rt*, |*s*|*t* » 1. Figure 3.C shows that this expression is a good approximation of the survival probability of the branching process. Since *β <* 1 when *s <* 0, the block is eventually lost. Equation (3.2) should be compared with the much faster decay of the survival probability for a single allele with fitness effect *sx* for *s <* 0, namely *p*_*t*_(*x*) −2*sxe*^−*sxt*^*/η*^2^.

**Figure 3:**
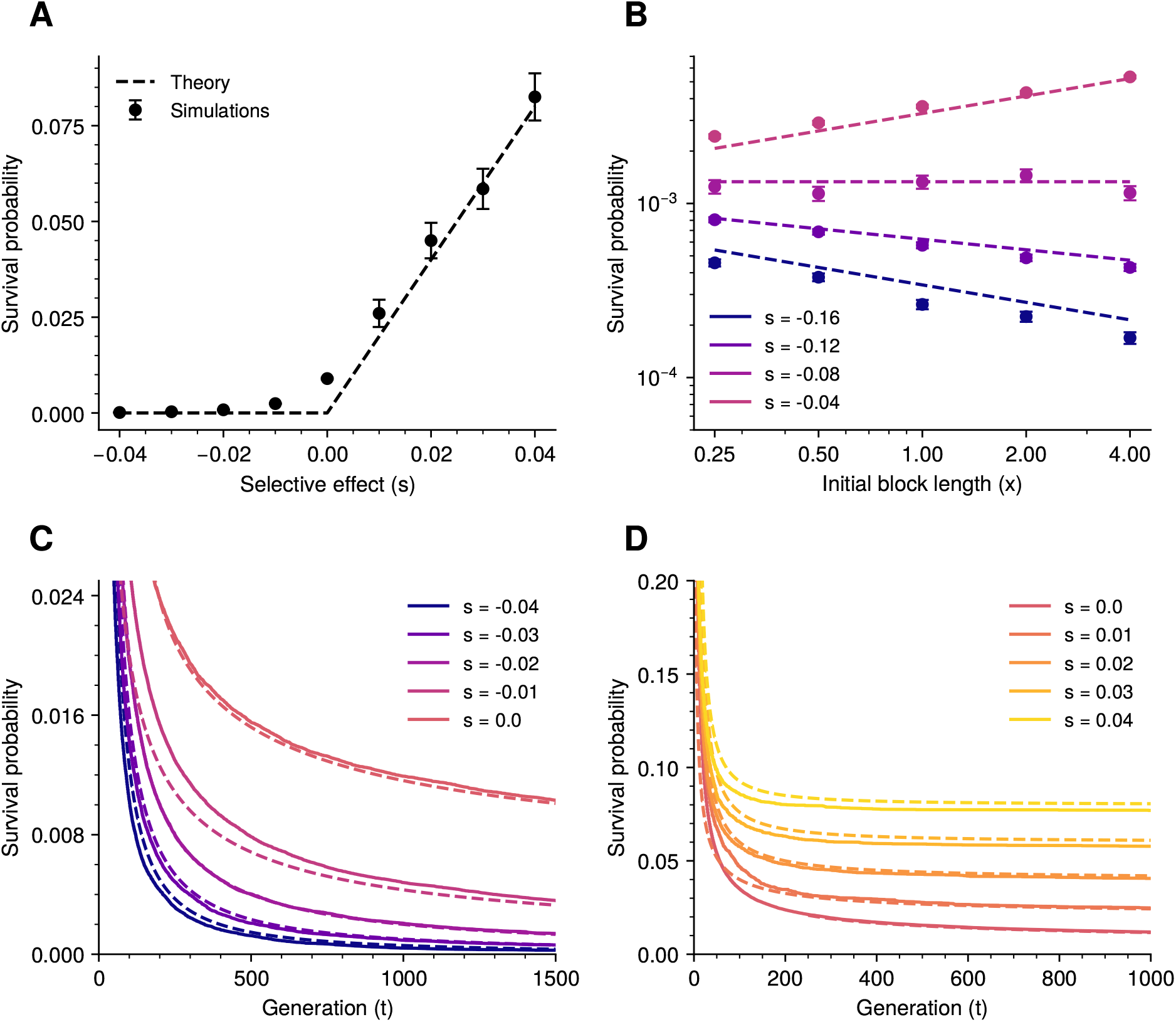
Survival probability of some portion of the introduced block. For all simulations *r* = 0.02 and for panels A, C, and D the initial block length is *x* = 1. (A) Probability of surviving *t* = 2,500 generations. Error bars indicate the standard deviation. The dashed line gives the probability that the process survives forever, obtained by letting *t* → ∞ in (3.2), (3.3), and (3.4). (B) Probability of surviving *t* = 1,500 generations, as a function of the initial block length. The dashed lines correspond to equation (3.2). (C) Survival probability as a function of time for negative selection. The dashed lines correspond to equation (3.2). (D) Survival probability as a function of time for positive selection. The dashed lines correspond to equation (3.4).

It is interesting to note that, if *r >* − *s* and so *β >* 0, equation (3.2) indicates that a larger block has a higher probability of leaving genetic descendants, even though this larger initial block is more deleterious. This counter-intuitive observation can be explained as follows. An individual carrying a subblock of length *y* has on average approximately 1 + (*r* + *s*)*y* children that also carry some part of that block. Recombination generates an additional *ry* net mean number of offspring, albeit with smaller block length. When *r >* − *s*, this additional number of offspring is sufficient to compensate the reduction in fitness induced by the block, and 1 +(*r* + *s*)*y* increases with the block length *y*. Conversely, when *r <* − *s* (and so *β <* 0), recombination is not strong enough to compensate for the additional negative selection, and the probability that some material survives decays with the length of the initial block. When *r* = −*s* and so *β* = 0, recombination and selection balance out and equation (3.2) reduces to the well-known expression for the probability of survival of a neutral locus, even though there is negative selection. These observations are illustrated in Figure 3.B.

#### Neutrality (*s* = 0)

If the introduced block is neutral (*s* = 0 and so *β* = 1), the survival probability decays as

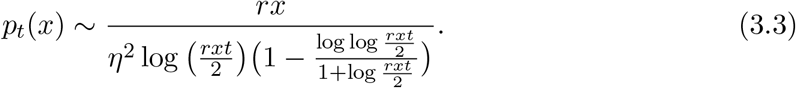

This result was already derived and discussed in Baird et al. (2003), see equation (7) there. Again, in comparison with a single allele (*r* = 0 and *p*_*t*_(*x*) ∼ 2*/*(*η*^2^*t*)) the survival probability decays much more slowly. It is quite likely that some of the introduced genetic material remains present, even after a very long time.

#### Positive selection (*s >* 0)

Finally, when selection is positive (*s >* 0 and so 1*/β <* 1), we compute in Section A.2 that

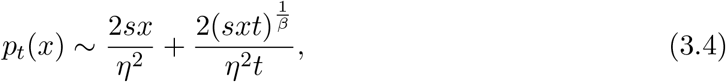

where once again we recall that in using the asymptotics of the diffusion approximation we are assuming that *r, s* « 1 and *rt, st* » 1. This expression is compared to simulations of the branching process in Figure 3.D, which shows that it is a good approximation. Since *p*_*t*_(*x*) has a positive limit 2*sx/η*^2^, there is a positive probability that some of the block persists forever (in the branching process) and so we can expect it to eventually fix in the population. A remarkable feature of this limit is that it does not depend on *r*. In particular, it remains valid in the absence of recombination (*r* = 0) and we recover as a special case Haldane’s formula for the fixation probability of a beneficial mutant. At the end of Appendix A.2 we outline a (mathematical) heuristic explanation, but at first sight this point is rather surprising. One might have expected that, by breaking the original block into pieces with smaller selective advantage, recombination counteracts selection and reduces the probability of fixation. Our result indicates that this effect is not visible at first order and, since we are only asking for survival of some part of the block, it is actually reversed at second order: the survival probability only decays to its limiting value polynomially in time in the presence of recombination, whereas it does so exponentially fast when *r* = 0 (*p*_*t*_(*x*) ∼ (1 + *e*^−*sxt*^)2*sx/η*^2^).

### 3.3 Probability of survival of a distinguished locus

The results in the last section concerned the probability that any subblock of the introgressed block survives to large times; they do not say anything about the location of surviving blocks. It is natural to ask whether there will be a block of introgressed genome around a particular locus at large times. Evidently this will be smaller than the probabilities calculated in the previous section, as only half of the offspring created through recombination will carry the distinguished locus.

Write *p*_*t*_(*x*_1_, *x*_2_) for the probability that a locus that initially has length *x*_1_ of the introduced block to its left, and length *x*_2_ to its right, survives until time *t*. Using the diffusion approximation we find that *p*_*t*_(*x*_1_, *x*_2_) can be approximated by the solution to

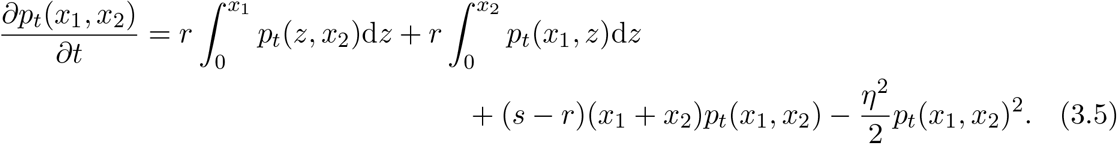

Consider first the case *x*_1_ = 0, so that *p*_*t*_(0, *x*) is the probability of survival of the left endpoint of the block. Whether blocks including this endpoint persist for arbitrarily large times is determined by the sign of *s* − *r/*2. Using methods entirely analogous to those used to prove the results of Section 3.2, writing 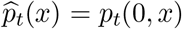 we show in Section A.3 that if *s > r/*2, then

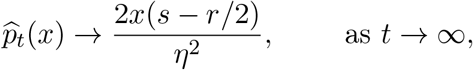

while if *s < r/*2,

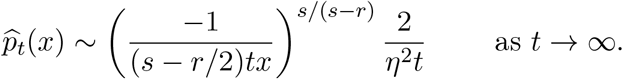

Notice that this is decaying to zero faster than the corresponding probability for a neutral locus. We were unable to find a closed form expression for the solution to equation (3.5) when both *x*_1_ and *x*_2_ are nonzero. However, it can be solved numerically. As we see in Figure 4, the theory matches the simulations well, although it somewhat overestimates the survival probability when *s/r* is large (when we expect the diffusion approximation to break down). We also see that there is some variability in survival probability along the block; if *s >* 0, then loci in the middle of the block have a somewhat higher chance of survival; if *s <* 0 it is lower. To understand why, note that although during a recombination event, the probability that the distinguished locus survives is 1*/*2, the probability that the recombination falls in the block to the left of the distinguished locus is *x*_1_*/*(*x*_1_+*x*_2_), in which case we expect a block of length with mean *x*_1_*/*2 to be ‘shed,’ and similarly with probability *x*_2_*/*(*x*_1_ + *x*_2_) the block that is shed has expected length *x*_2_*/*2. Overall, if a recombination takes place, the amount of block shed has mean 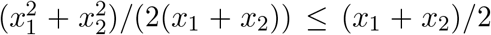. Recombination events that preserve a distinguished locus in the interior of the block are less effective at shedding introduced block than those preserving the left endpoint. In the case 2*s > r* we find a lower bound on the amount by which the survival probability of a fixed locus at an interior point of the block exceeds that of the locus at an endpoint in Section A.3.

**Figure 4:**
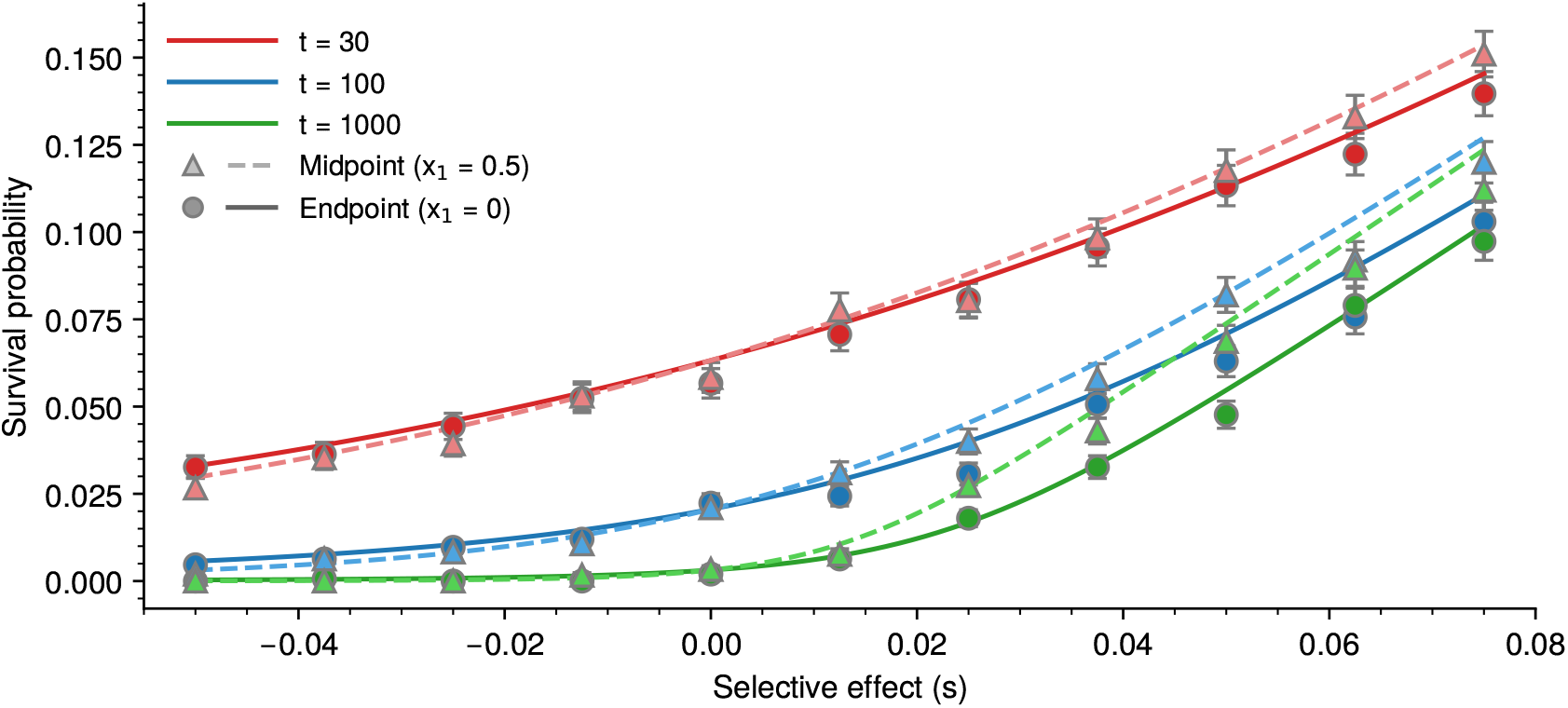
Probability of survival of a distinguished locus. The curves show the predicted probability that a distinguished locus will survive to times 30, 100, 1000 (red, blue, green), and the points show means ±1 standard deviations for simulations with 3000 replicates; *r* = 0.05, *x* = 1. Solid lines and circles are for a locus at the end; dashed lines and triangles for a locus at the midpoint.

### 3.4 Distribution of introgressed block lengths (*s > r*)

The results of Section 3.2 give the probability that at least some genetic material survives, regardless of the amount of that material. We now turn to the typical length and number of copies of the surviving blocks. Let *M*_*t*_(*x, y*) be the mean number of blocks of length at least *y* in generation *t*, when the population starts from an initial block with length *x*, and write *M*_*t*_(*x*) for lim_*y*↓0_ *M*_*t*_(*x, y*), the mean total number of blocks. In Section A.5, we obtain an approximation for these quantities by studying the Fokker–Planck equation solved by the mean block length distribution of the diffusion approximation. For *s* ≠ *r* we show that

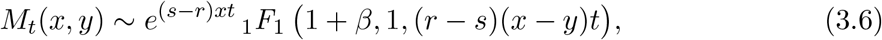

where the confluent hypergeometric function _1_*F*_1_ is defined by

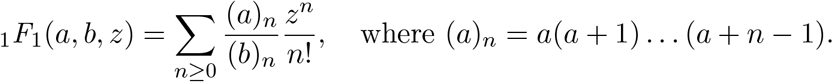

When *b > a*, it has the simpler integral representation

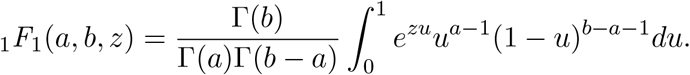

The asymptotic behaviour of this function is well known:

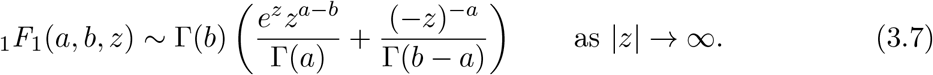

The behaviour of our approximation depends strongly on whether *s > r*, in which case the initial block can persist for a long time; or *s < r*, in which case blocks are rapidly lost. We start with the former case.

When *s > r* (and so *β <* − 1), applying (3.7) to (3.6), the second term dominates and we find

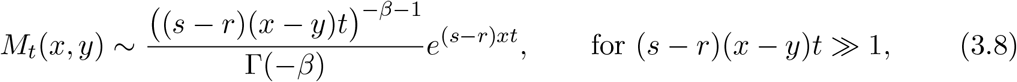

and the limiting cumulative distribution of block lengths is given by

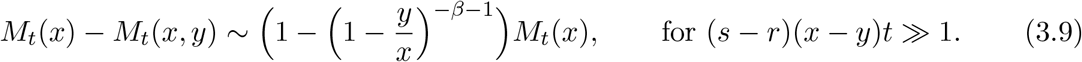

The expected number of copies of the initial block grows approximately as *e*^(*s*−*r*)*xt*^, whereas, from equation (3.8), the expected total number of blocks, *M*_*t*_(*x*), grows as ((*s* − *r*)*xt*)^−*β*−1^*e*^(*s*−*r*)*xt*^ (with −*β >* 1). Thus, even though the initial block can persist at large times with positive probability, its frequency among surviving blocks vanishes as *t* → ∞ (due to the factor *t*^−*β*−1^). Nonetheless, equation (3.9) indicates that the blocks that survive remain of comparable length to the introduced block, and that their lengths are asymptotically Beta(1, −*β* − 1) distributed over the original block [0, *x*]. The distribution of block lengths is therefore tilted towards smaller blocks if *s <* 3*r* (and so −*β <* 2) and towards larger blocks if *s >* 3*r* (when −*β >* 2). Moreover, the distribution of blocks concentrates on a Dirac mass at 0 if *s* → *r* (and so *β* → −∞), indicating that in that case block lengths become of a smaller order than the initial block. Conversely, the distribution concentrates on a Dirac mass at *x*, the length of the introduced block, if *s* » *r* (and so *β* → − 1). In this case, the initial block behaves almost as a single allele under positive selection, without recombination, and setting *β* = −1 in equation (3.8) we recover that the mean number of blocks grows exponentially at rate *sx*.

We have provided expressions for the expected numbers of blocks with lengths in a specified interval at large times, but we could equally have reported *m*_*t*_(*x, y*), the corresponding expected density of blocks of length *y < x* (so that 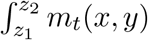)d*y* gives the mean number of blocks with lengths in the interval [*z*, *z*]). Differentiating *M*_t_ (*x, y*) we find

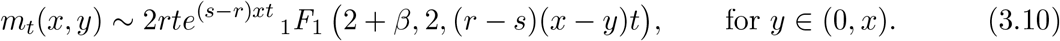

Because there is a positive probability of the original block surviving intact, there will be a delta-function at *y* = *x*. Our approximations for *m*_*t*_(*x, y*) and *M*_*t*_(*x*) − *M*_*t*_(*x, y*) are illustrated numerically in Figure 5.A-B. Overall, the theoretical prediction for the distribution of block lengths matches the simulations well.

**Figure 5:**
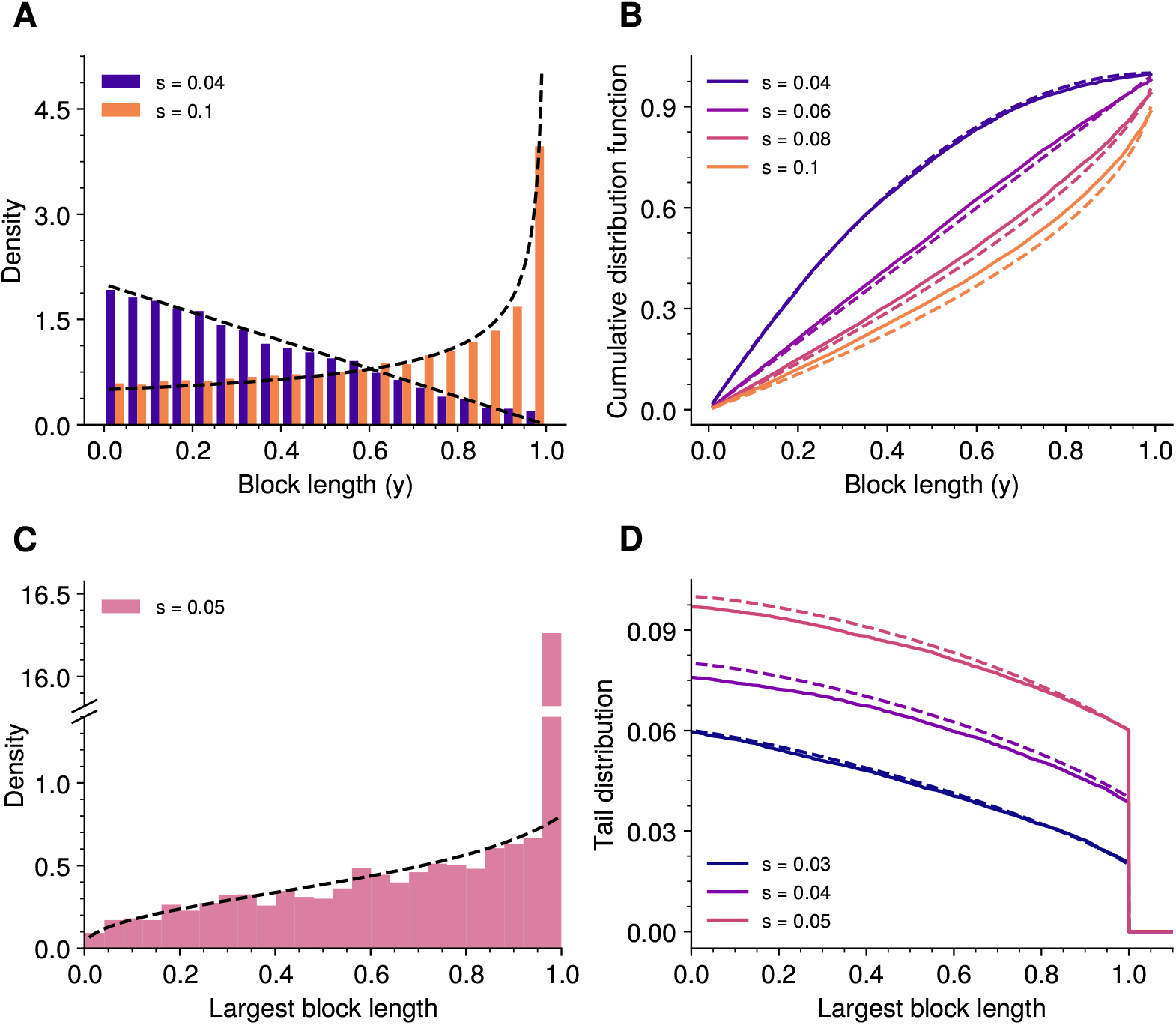
Distribution of block length, *s > r*. In all simulations *r* = 0.02 and *x* = 1. Distribution (A) and cumulative distribution function (B) of block lengths in one replicate at time *t* = 2,500 for one replicate. Simulations are culled at 10^5^ blocks as described in Section 2.5; dashed curves correspond to (3.9). Distribution (C) and tail distribution function (i.e. probability length is greater than *y*, D) of the length of the largest block in the population, for 10^5^ replicates for each value of *s*. Simulations are conditioned on survival of the population, and stopped either after *t* = 2,500 generations or upon reaching 5 × 10^5^ blocks. Dashed lines correspond to (3.11).

The results above indicate that when *s > r*, large blocks can persist at large times. We can also derive the distribution of the longest block present in the population after *t* generations. Let 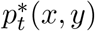 be the probability that the largest block in the population in generation *t* is larger than *y*, given that the population started with an initial block of length *x*. By studying the generating function of the diffusion approximation, we show in Section A.4 that

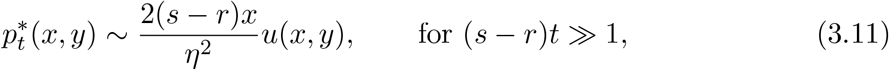

where *u*(*x, y*) solves

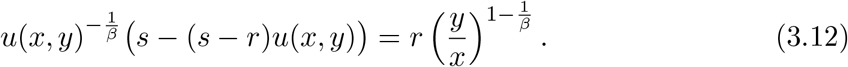

This equation does not have a unique solution, but *u*(*x, y*) is uniquely specified by taking the solution with values in [1, ∞). Even then, there is no closed form for *u*(*x, y*), but as we see from the numerical solution in Figure 5.C-D it provides a good approximation to the simulations. The initial block (of length *x* = 1) has a positive probability of survival, as indicated by the atom at *y* = 1 in Figure 5.C, and by the jumps in Figure 5.D. Since this initial block cannot be created by recombination from larger blocks, it behaves as a single allele with fitness (*s* − *r*)*x* and by Haldane’s formula the probability that an intact copy survives is 2(*s* − *r*)*x/η*^2^. (We note that this is consistent with (3.11), since *u*(*x, x*) = 1 is a solution of (3.12) when *y* = *x*.) Since equation (3.4) shows that the probability that the population survives is 2*sx/η*^2^, we deduce that the probability that the largest block is strictly smaller than the original one is 2*rx/η*^2^. In this case, the distribution of the largest block is not strongly dependent on *s*, as can be seen from the similarity of the curves in Figure 5.D.

### 3.5 Distribution of introgressed block lengths (*s* ≤ *r*)

We now consider the regime *s* ≤ *r*, in which large blocks eventually go extinct.

**Case *s < r***. In the case *s < r*, using (3.7) in (3.6), it is the first term that dominates, and we find

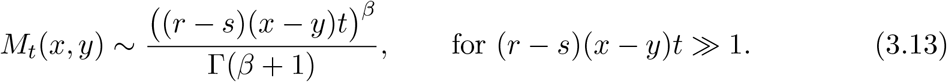

In this regime, surviving blocks are typically of length 1*/*(*rt*). For *y* « *x*, we compute that

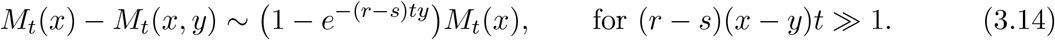

Equation (3.14) suggests that surviving blocks are, on average, asymptotically exponentially distributed with mean 1*/*((*r* − *s*)*t*). In this regime, large blocks are lost rapidly through recombination, which is the main driving force of the system. The limiting exponential distribution can be understood intuitively. Suppose first that our introgressing block were neutral, as in Baird et al. (2003). At large times, if we record all the crossovers that have affected a block of genome of length *x* (corresponding to the position of our introgressing block) in an individual at time *t*, then scanning from left to right across the block they (approximately) follow a Poisson point process with rate *rt*. The typical distance between crossovers, and so the length of an introgressed block (if one exists), is therefore distributed as an exponential random variable with mean 1*/*(*rt*). Our result shows that this heuristic is only valid if the introgressed material is neutral. Otherwise, selection makes the blocks on average longer when *s >* 0 or shorter when *s <* 0, as one expects. If the recombination rate is known, the selective effect could (in principle) be estimated from our expressions by computing the mean length of introgressed blocks.

Equation (3.13) can be combined with the expressions for the survival probability to estimate the mean number of blocks conditional on survival. The result depends on whether the selective effect of the block is negative, in which case

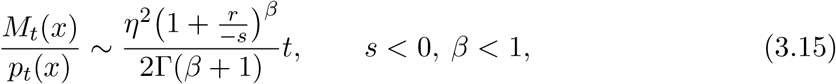

neutral, for which

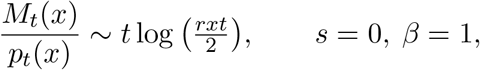

or positive, and

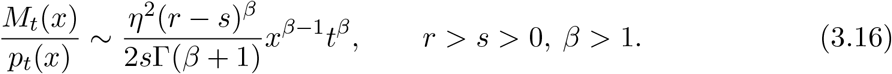

When selection is negative, the number of blocks grows linearly with time during a successful introgression. This is characteristic of critical branching processes, which correspond to the early dynamics of a neutral allele reaching fixation. This indicates that, typically, the introduced block is very rapidly broken down into small pieces, after which the resulting blocks behave almost as neutral loci. We saw in equation (3.2) that a larger initial block has a higher or lower probability of surviving depending on whether −*s < r* or −*s > r*. The right-hand side of equation (3.15) does not depend on *x*, which indicates that the length of the initial block does not influence the number of blocks conditional on survival. These observations remain almost valid in the neutral case (*s* = 0) up to logarithmic terms, and have already been derived and discussed in Baird et al. (2003).

On the other hand, when selection is positive (0 *< s < r* and so *β >* 1) the number of blocks that survive grows as ((*r* − *s*)*t*)^*β*^*x*^*β*−1^. Note that this is a power law growth, slower than an exponential (for 1 *< β <* ∞).

These results are illustrated for positive selection (*r > s >* 0) in Figure 6. This realisation fulfils both our predictions, that the average block length decays approximately as 1*/*((*r* − *s*)*t*) when *t* is large, and that the block length distribution becomes asymptotically exponential. However, in simulations we also observe a second type of possible behaviour that does not match our predictions. One example of such a realisation is displayed in Figure S5 in which the distribution is dominated by several atoms, corresponding to clones of the same block. We believe that such realisations are characterised by a rapid loss of large blocks in the early stages of the simulation; surviving blocks at large times are confined to a small region of the genome, more reminiscent of the behaviour we see in the neutral case.

**Figure 6:**
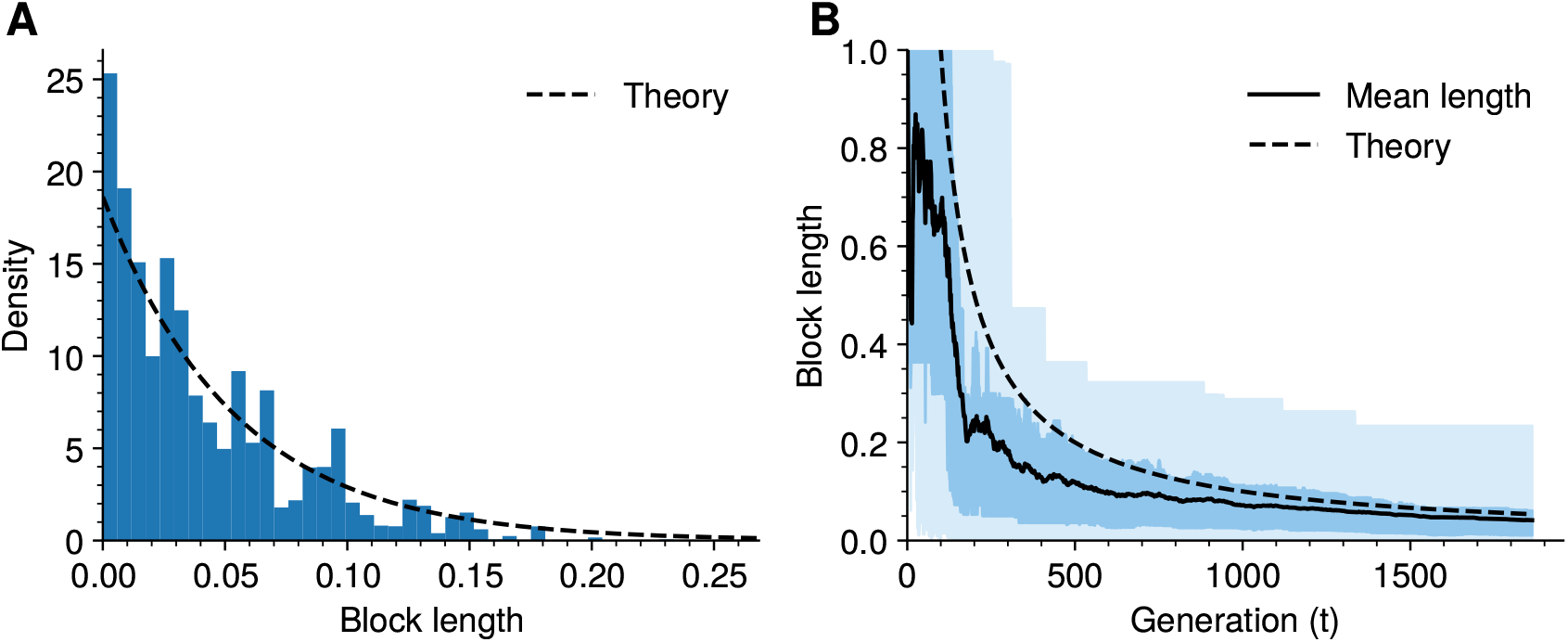
Distribution of block lengths, *s > r*. (A) Distribution of block lengths at generation *t* = 2,500 in one realisation of the process, conditioned on survival and with *s* = 0.01, *r* = 0.02, and *x* = 1. (B) Length of the blocks through time for the same realisation. The solid line shows the average block length in the population; the region with a lighter shade represents the largest/smallest blocks; and that with a darker shade the 25% and 75% quantiles of the distribution of block lengths. Dashed lines are obtained from (3.14).

#### Case *s* = *r*

Lastly, for the sake of completeness, we provide the limit of the mean number of blocks when *r* = *s*. We show in Section A.5 that in this case

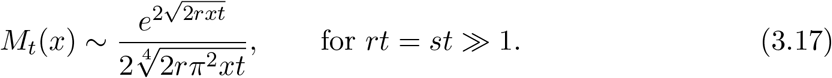

Moreover, for *y* « *x*,

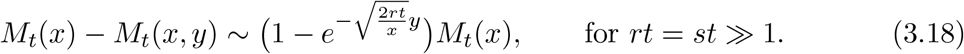

The situation is intermediate between the other two cases. Block lengths no longer remain macroscopic, but decay as 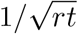. The expected number of blocks grows as a stretched exponential, again intermediate between the exponential and polynomial growths observed in the other two cases.

### 3.6 Total mass of introgressed material through time

From *m*_*t*_(*x*, d*y*), we can estimate the total expected amount of introgressed material as a function of time. Substituting from equation (3.10), when *s* ≠ *r*

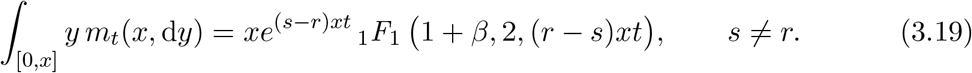

The corresponding expression for *s* = *r* can be found in (A.33). In Figure 7 we compare this prediction to simulations for different strengths of (positive) selection relative to map length. We also compare to the case with no recombination. When selection is weaker than recombination, mass increases more slowly than exponentially, because blocks become ever smaller. When selection is stronger than recombination, the mass increases approximately exponentially, at a rate intermediate between the rate with no recombination and the contribution from intact blocks. Note that the proportion of intact blocks decreases over time.

**Figure 7:**
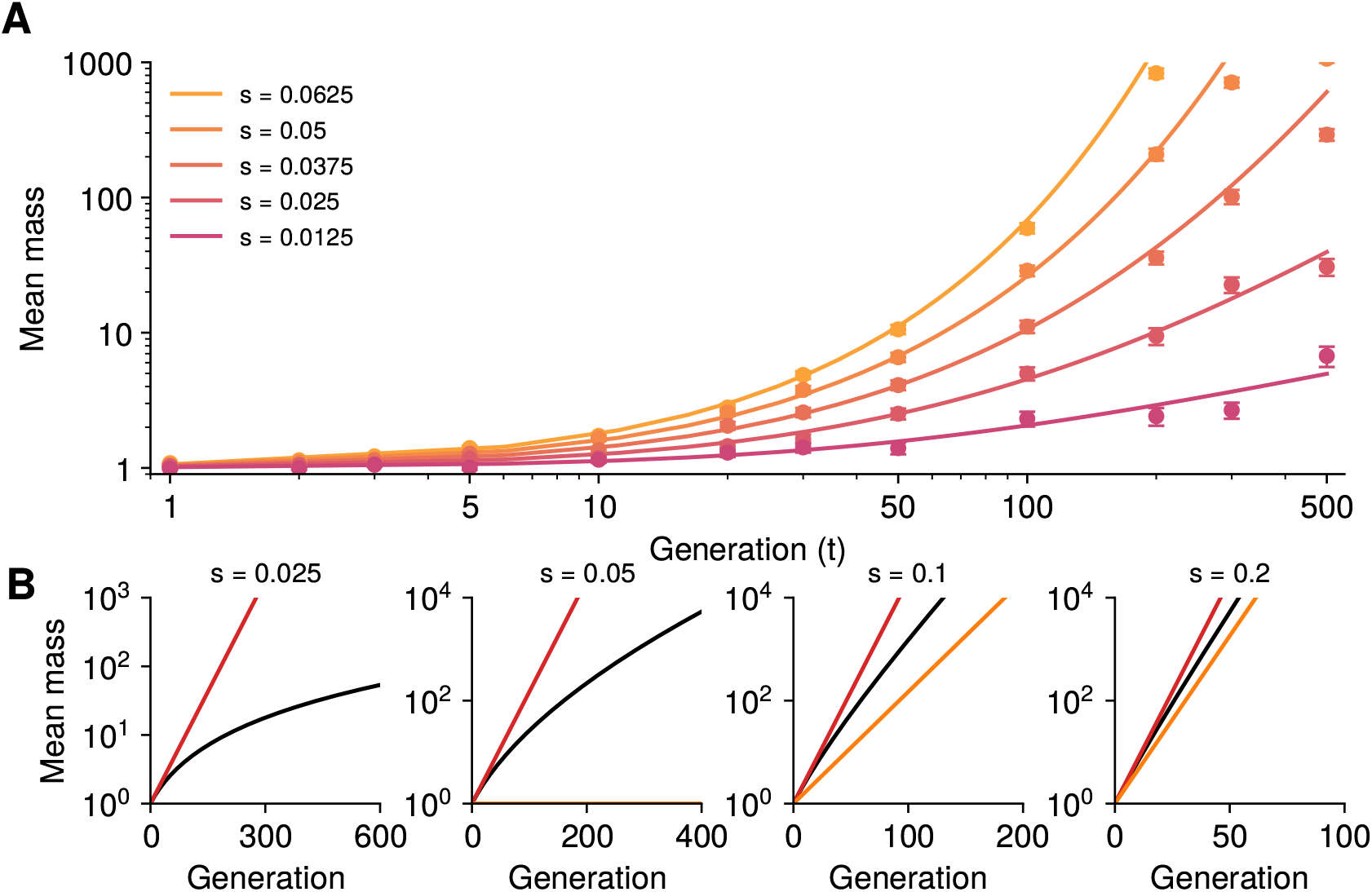
Increase in total mass of blocks through time. (A) Increase in total mass of introgressed material through time for *r* = 0.05, *x* = 1. This agrees well with the theory (solid lines). Three thousand replicates were run until loss or more than 10^5^ blocks. (B) Comparison of increase in mean mass (black) with the increase with no recombination (red), and with the contribution from the intact block (orange).

### 3.7 Coalescence times and location of blocks on the genome

It is also interesting to examine the time at which two individuals, sampled at random from among those currently carrying some portion of the introduced block, had their most recent common ancestor, that is their coalescence time. The blocks that the individuals carry must be subblocks of the block carried by their common ancestor. Knowing coalescence times will enable us to make predictions about the length of the block carried by the ancestor of a typical pair of individuals, which in turn tells us about the maximum possible distance on the chromosome between the blocks carried by the sampled individuals.

There is no simple expression for the distribution of the coalescence time of two uniformly chosen individuals in the branching population, even under the diffusion approximation. Instead, we will consider *C*_*t*_(*x, κ*), the expected number of pairs of individuals in generation *t* for which the most recent common ancestor lived before generation *κ* (starting from an initial block of length *x*), divided by *C*_*t*_(*x, t*) (the expected *total* number of pairs of blocks in the population at time *t*). Although there is no simple relationship between this quantity and the probability that a randomly chosen pair coalesced before time *κ*, we expect it to scale in a similar way. Note that, unlike in coalescent theory, time *κ* is going forwards and not backwards here.

We consider first the regime *s < r*, in which block lengths vanish asymptotically. The typical time at which coalescences occur depends on whether selection is positive, neutral, or negative. All computations are carried out in Section A.6. The results are illustrated in Figure 8.

**Figure 8:**
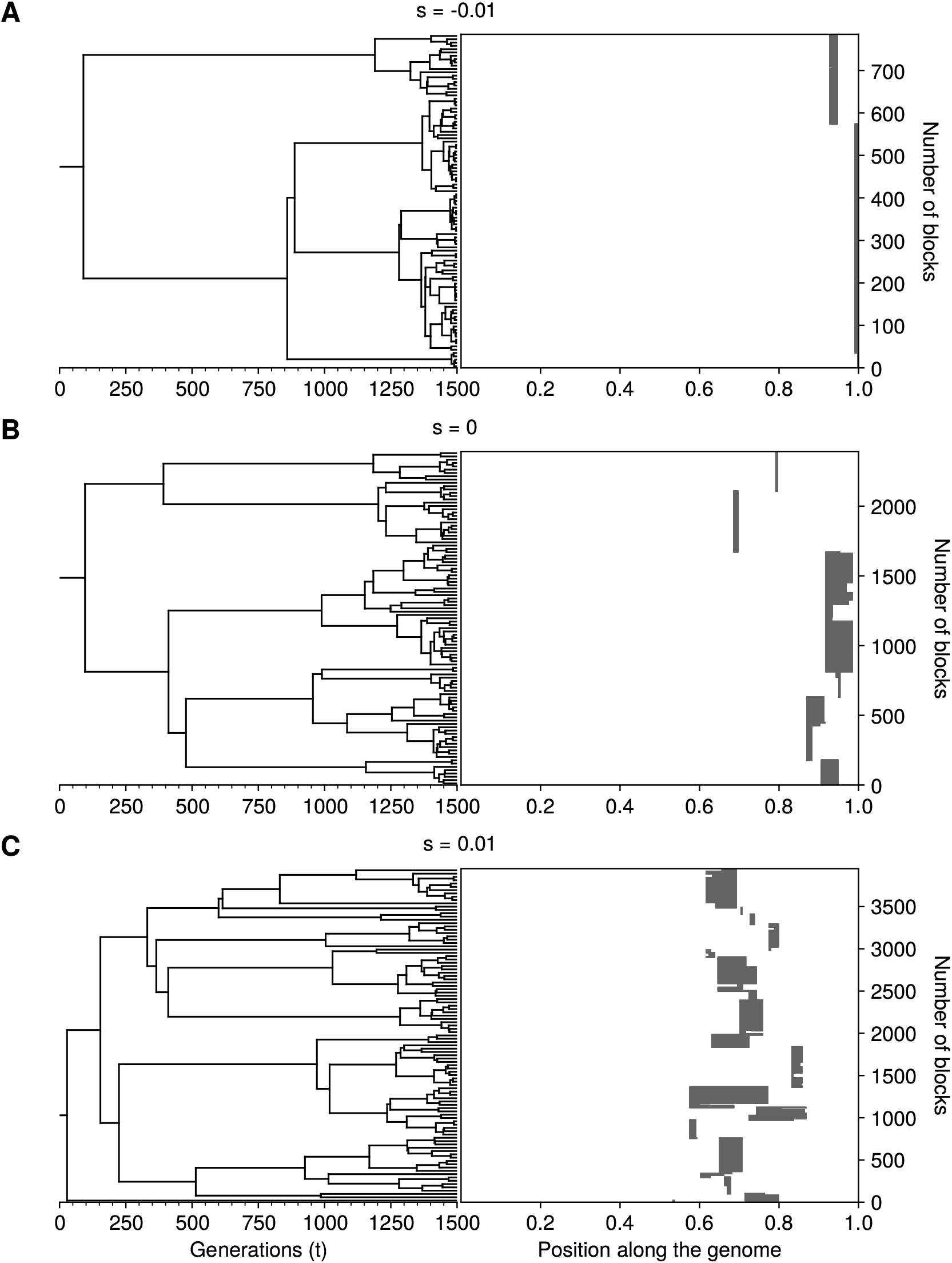
Example of genealogies. Three independent realisations of the branching process at generation *t* = 1,500, conditioned on survival and with *r* = 0.02, *x* = 1 and *s* = −0.01, 0, and 0.01 in (A), (B), and (C) respectively. The panel on the right hand side shows the state of the population at the final generation, and the panel on the left shows the genealogy of a sample of 100 individuals.

#### Weakly positive selection (0 *< s < r*)

If 0 *< s < r* (and so *β >* 1) we find that

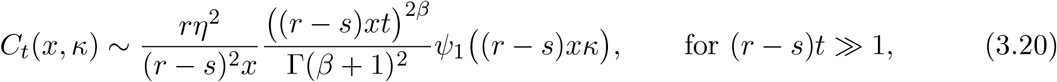

where *ψ*_1_ only depends on *β* and is defined in (A.37). It is increasing and satisfies

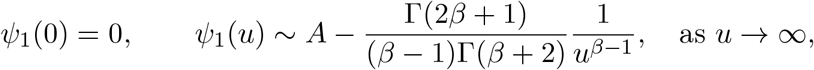

for some constant *A*.

Substituting into (3.20),

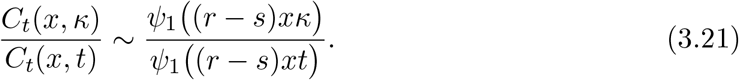

The form of *ψ*_1_ implies that this will start to be significantly different from 1 at times *τ* of order 1*/*(*x*(*r* − *s*)). At such times, ancestors carry blocks of comparable length to the length *x* of the introduced block. Therefore, even though (as we see from equation (3.14)) most surviving blocks will be small compared to *x*, they are found across a genomic region of comparable length to it. These results are illustrated by the realisation in Figure 8.C, where branch points occur early on, and surviving blocks span half the initial block. Early coalescence times are expected under supercritical branching (corresponding to positive selection), but whereas for a single positively selected locus the time to the most recent common ancestor of two randomly chosen individuals would have an exponential tail, here we see that the ratio in (3.21) approaches one at polynomial speed as (*r* − *s*)*xκ* increases, indicating that the limiting distribution of branch times has a heavy tail.

#### Negative selection (*s <* 0)

If *s <* 0 (and so *β <* 1),

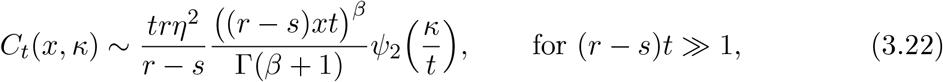

for another another increasing function *ψ*_2_ defined in (A.38) that only depends on *β* and satisfies

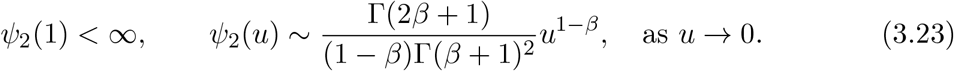

The situation is very different from positive selection. As above, we calculate the ratio *C*_*t*_(*x, κ*)*/C*_*t*_(*x, t*). In this case, as *κ/t* → 0 it is equivalent to ∼ (*κ/t*)^1−*β*^ (with *β <* 1), and so we expect that the most recent common ancestor of two individuals carrying blocks of introgressed material in generation *t* will have lived at a time, *τ*, of the same order as *t*. This is reminiscent of critical branching processes and is in line with the intuition that during a successful introgression of deleterious material, the blocks rapidly become very short and behave almost as neutral loci. Moreover, since the mean block length in generation *τ* decays as 1*/*((*r* − *s*)*τ*), the most recent common ancestor can be expected to carry a block with length of the same order as those at time *t*, that is 1*/*(*r* − *s*)*t*. As a result, we expect that all surviving blocks are found in a very narrow region of the genome, which is of comparable length to that of the blocks themselves. We see these two predictions reflected in the realisation shown in Figure 8.A. We saw in Section 3.2 that if *r >* −*s >* 0, recombination acts to compensate for some of the negative selection and impacts survival probabilities; here we see that it also impacts the genealogy of the population: since *C*_*t*_(*x, τ*)*/C*_*t*_(*x, t*) ∼ (*τ/t*)^1−*β*^ for *τ/t* « 1, coalescence times are deeper if *r >* −*s* (0 *< β <* 1) than if *r <* −*s* (*β <* 0).

#### Neutrality (*s* = 0)

In the neutral case (*s* = 0), we show that for *v* ∈ (0, 1),

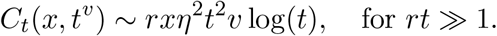

Setting *v* = 1, the total number of pairs in this case grows as *rxt*^2^ log(*t*). Thus, coalescences typically occur at generations *κ* for which log(*κ*) is of the same order as log(*t*), that is, *κ* = *t*^*u*^ for some *u* ∈ (0, 1). At that time, individuals have blocks of length 1*/rt*^*u*^ « 1. This length is negligible compared to the length of the initial block, and surviving blocks are spread across a small genomic region (compared to the length of the introduced block), but which remains much larger than their typical length of 1*/*(*rt*). See Figure 8.B for a numerical illustration. A full description of the genealogy in this case can be found in Foutel-Rodier and Schertzer (2023).

#### Strongly positive selection (*s > r*)

Finally, when *s > r*, we prove in Section A.6 that

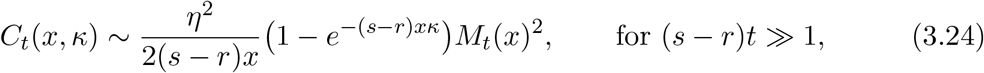

where *M*_*t*_(*x*) is given by equation (3.8).

Similarly to the regime 0 *< s < r*, coalescence times are of the order 1*/*((*s* − *r*)*x*) and occur close to the origin. Note however that the limiting distribution of branch times implied by (3.24) has an exponential tail, as opposed to the polynomial tail of (3.21) when 0 *< s < r*, indicating that coalescences occur even closer to the origin.

### 3.8 Beneficial locus in a deleterious background

We turn to the situation described in Section 2.4, in which the initial block contains many alleles with a negative impact on the trait and one point locus with a beneficial allele. We record a block as a pair (*y*_1_, *y*_2_) giving the length of deleterious material to the left and right of the beneficial allele, respectively. The population starts from an initial block (*x*_1_, *x*_2_) with total length *x* = *x*_1_ + *x*_2_. Clearly the process has a positive probability of surviving forever, which can be approximated as the stationary solution to

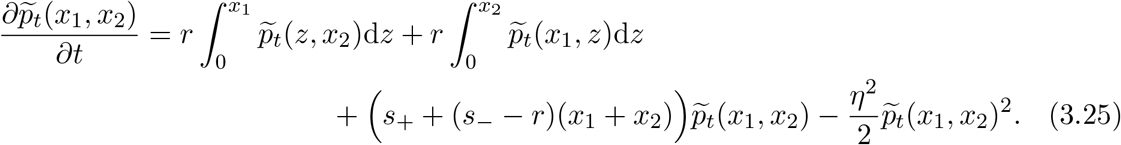

We could not find a closed-form expression for the stationary solution to this equation, although a comparison of its numerical solution to simulations is shown in the top panel of Figure 9. Instead, we focus on computing the amount of deleterious material which hitchhikes with the beneficial locus. Let us denote by 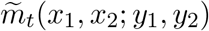 the mean density of blocks (*y*_1_, *y*_2_) at generation *t*, and by 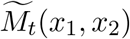 the mean total number of blocks. Let

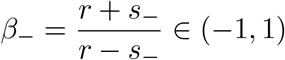

be the characteristic exponent associated with the deleterious material. In Section A.7, we show that, for *t* » 1,

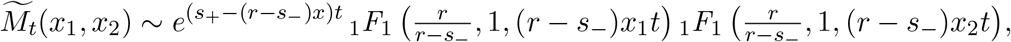

and

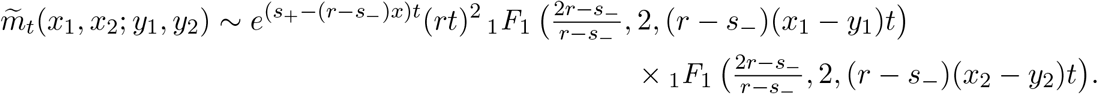

**Figure 9:**
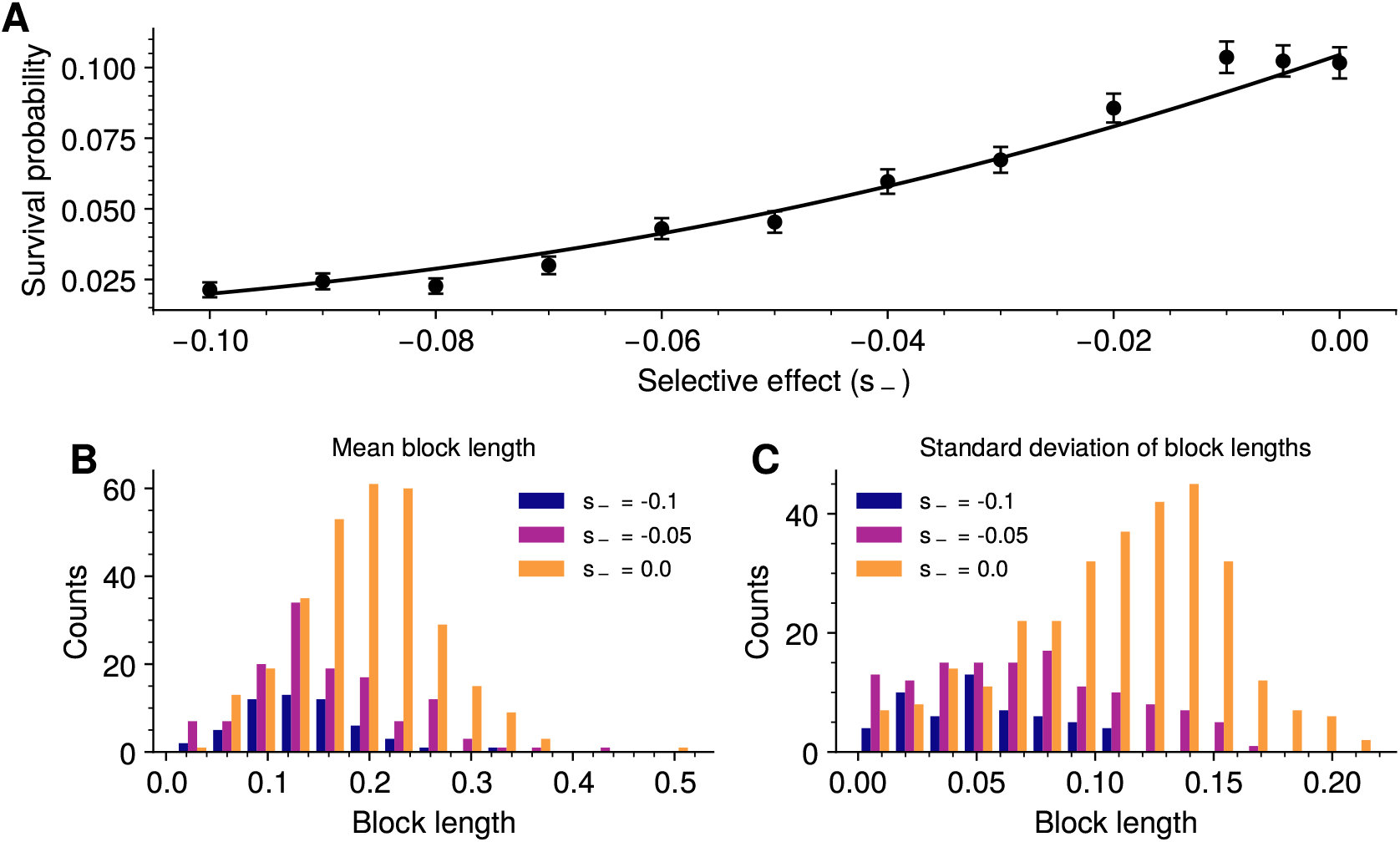
A beneficial allele embedded in a deleterious block. (A) Probability of survival of a discrete favourable allele as a function of *s*_−_, for *r* = 0.1, *x* = 0.1. The selectively advantageous allele, with *s*_+_ = 0.05, is embedded at the middle of the block. There are 3,000 replicates and the simulation stops when there are more than 10^5^ blocks. The solid line is the theoretical prediction (obtained as the numerical solution to equation (3.25)). (B, C) Distribution of the mean and standard deviation of block length across replicates for different background selection coefficients. This is the distribution of blocks after 100 generations in realisations in which the total number of blocks eventually reaches 10^5^ blocks.

These expressions can be further simplified by assuming that *rt* » 1,

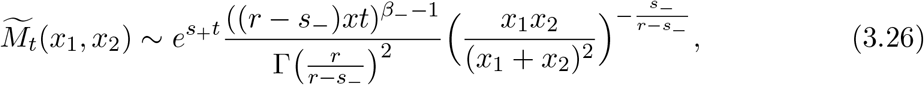

and

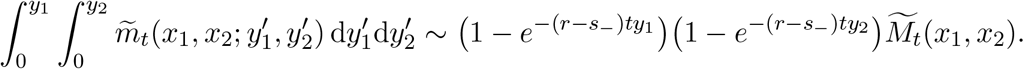

If the deleterious material were absent, the mean number of copies of the beneficial allele would grow as 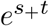. This growth is reduced by a factor 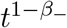 due to the fitness loss incurred by the linked deleterious material. Moreover, as in the absence of the beneficial allele, recombination breaks down the blocks to a typical length of 1*/*((*r s*)*t*) at time *t* and the lengths of the part of the block on each side are independent and exponentially distributed. Again, if the recombination rate *r* is known, one could, in principle, estimate the selective effect of the deleterious material from the mean length of introgressed blocks. Note also that (3.26) is maximal when *x*_1_ = *x*_2_ = *x/*2, that is when the locus is initially located at the centre of the block. This mirrors what we saw in Section 3.3; recombination is most effective at removing the deleterious material when it is equally spread on both sides of the beneficial locus.

Finally, during the introgression of the beneficial allele, some purely deleterious blocks are generated which do not contain the beneficial locus. Let us denote by 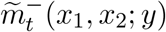, the mean density of such blocks with length with length *y* at time *t*, starting (in the notation above) from a block (*x*_1_, *x*_2_) containing the beneficial allele. At the end of Section A.7 we show that

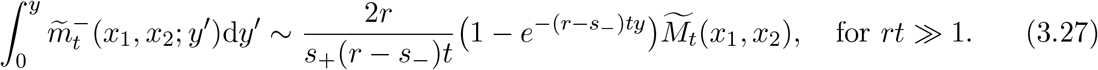

The first implication of this expression is that the mean number of purely deleterious blocks grows more slowly than the mean number of blocks containing the beneficial allele, by a factor of 1*/*((*r* − *s*_−_)*t*). Since deleterious blocks are rapidly lost, the only such blocks present in the population at large times are those that were formed recently, by recombination from a block containing the beneficial allele. Because the latter blocks are of length of order 1*/*((*r* − *s*_−_)*t*), the rate at which they experience crossovers is of the same order, which explains why there are fewer deleterious blocks. The second conclusion one can draw from equation (3.27) is that the length of an average purely deleterious block is exponentially distributed with mean 1*/*((*r* − *s*_−_)*t*). This can be understood intuitively by noting that the length of a typical block containing the beneficial allele is the sum of two exponential random variables. Splitting such an interval at a uniform crossover yields an exponential random variable.

These results are illustrated in Figure 9. The theoretical prediction for the survival probability, obtained by solving equation (3.25) numerically, matches the simulations well. The lower panel plots the distribution of block lengths that have been carried along by the beneficial locus after 100 generations, where only realisations of the simulation in which the total number of blocks eventually reaches 10^5^ are included. Selection against the deleterious background reduces the mean size of the blocks relative to the neutral case (shown in yellow), but there is substantial overlap, which would make it impossible to detect linked selection. Stronger selection against surrounding genome would greatly reduce the probability of survival (as we see in the upper panel). In these simulations the beneficial allele reaches 10^5^ copies, and so can be expected to fix. Whilst the favoured allele is still rare, recombination will continue to reduce the sizes of the associated blocks. Once the favourable allele becomes common, these blocks will recombine with each other, and selection will reduce the amount of deleterious material until what fixes will be a small associated block. We do not consider this process here.

## 4 Discussion

We analyse the fate of a single block of genome that is introduced into a large homogeneous population, under the simple assumption that fitness is proportional to the amount of introduced material. Our analysis assumes a branching process, and so is restricted to blocks small enough that no more than one crossover is likely. After hybridisation, a long genome will suffer multiple crossovers, leaving descendants with multiple blocks. However, after a few generations, most descendants will carry only one block, after which our analysis applies. Baird et al. (2003, Section 6) analysed the case of multiple crossovers, assuming neutrality (or a fixed selective advantage), and derived the mean number of surviving blocks. However, other quantities were intractable. Thus (following Baird et al. (2003)) it seems simplest to simulate the first few generations, and then apply our results to the distribution of block lengths. Similarly, our analysis breaks down when blocks are so small that they contain just a few loci. Under a branching process, the fate of sets of discrete alleles can be readily calculated (Sachdeva and Barton, 2018b). Our analysis describes introgression from blocks of a few cM down to their disintegration into a few discrete loci.

The outcome of the introgression depends on the strength of selection relative to recombination, *s/r*. If the introduced material is deleterious (*s <* 0), it will ultimately be lost, and the chance that any material survives for *t* generations decays as 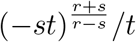 for *s <* 0; surviving material will be scattered into small blocks, with total mass ((*r* − *s*)*xt)* ^2*s*/(*r*™*s*)^. If the material is favourable, the chance that any descendants survive is 2*sx/η*^2^, exactly as for a single locus. However, the original block can only survive if selection is stronger than recombination (*s > r*); in that case, either the original block will survive (with probability 2(*s* − *r*)*x/η*^2^), or a large subblock will survive (with probability 2*rx/η*^2^). The largest surviving block will increase fastest, and will dominate the population; yet, it may be out-numbered by the descendant subblocks that it generates. In contrast, if selection is positive, but weaker than recombination (*r > s >* 0), the surviving blocks will become ever smaller (1*/*((*r v s*)*t*)), even as the total mass (and hence, mean fitness) increase as 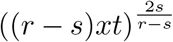. We also consider a simple extension, following the increase of a beneficial allele that is embedded in a block of deleterious genome; this will fix with positive probability, carrying with it deleterious genome with typical size 1*/*((*r* − *s*_−_)*t*) after *t* generations.

Since the size of surviving blocks depends strongly on the strength of selection relative to recombination, one can imagine inferring selection strength from block size. We consider two questions: first, the density of negative selection (*s*_−_ */r <* 0) around a beneficial allele, and second, whether selection is concentrated at one causal locus, or spread over some region of genome.

In the simplest case, a beneficial mutation will carry to fixation blocks of the original genome on either side, each exponentially distributed with mean map length 1*/τ*, where *τ* is the time taken to reach high frequency (Maynard Smith and Haigh, 1974; Barton, 1998). Selection against these accompanying alleles will reduce the size of these regions, and could be estimated if we knew the time *τ* during which recombination had acted; this might be possible using the method of Speidel et al. (2019), which depends on the shape of the genealogy at the focal locus. However, there is wide variation in the size of the surrounding block, making such estimates inaccurate (Figure 9). Also, deleterious alleles may be recessive, altering the dynamics. (Note that the time at low frequency may be much longer than the naive formula (1*/s*) log(4*N*_*e*_*s*) which assumes panmixis: population structure or fluctuations in selection may extend the time *τ*, so that selection would be severely underestimated.)

It may seem more feasible to find whether positive selection is concentrated at one locus, or is instead spread over some region. Predictions here are quite different: in the first case, one would see a ragged set of regions on either side, each with an exponential distribution, but always containing the discrete positively selected locus. In contrast, if an intact block arose, with net advantage greater than the maximum recombination rate between the positively selected loci (*s > r*^∗^), the whole region would fix, with a ragged neutral region surrounding it. In the first case, the region that sweeps to fixation has map length of order 1*/τ* = *s/* log(4*N*_*e*_*s*), whereas in the second, it has map length *r*^∗^+*s/* log(4*N*_*e*_*s*). Since we do not know *s* accurately, it may not be possible to distinguish these cases simply from these sizes. Rather, one would need to test whether the distribution of block lengths has either a mode at zero, or a strictly positive minimum value. One might do better with replicates, possible for experimental evolution or with independent origins of (say) pesticide resistance, although there might well be heterogeneity in which favourable regions are picked up.

The most obvious application of our result is to help understand the introgression of Neandertal genome into the modern human population. Eurasians carry around 2% of Neandertal-derived genome, in blocks of up to several hundred kb. These blocks are each typically rare (*<* 10%, say), but together cover much of the genome, in a heterogeneous distribution (Petr et al., 2019). The time course of introgression can be traced using ancient DNA, which shows that blocks have become smaller over time (Iasi et al. (2024) Fig.4c); the fraction of introgressed genome has decreased (though this may in part be because smaller blocks are less easily detected), but some blocks have increased in frequency. Assuming either neutrality, or selection on at most one locus in each block, their length decreases purely by recombination, allowing their time of origin to be inferred. This ‘recombination clock’ implies a single pulse of hybridisation, about 46,000 years ago, or around 1,600 generations, assuming a 28 year generation time (Iasi et al., 2024).

What do we expect to see, if selection acts in proportion to block length, as our model assumes? A single F1 individual will leave a random number of descendants, with a net reproductive value that is highly variable, and that may be inflated if the introgressing genome confers some net selective advantage (Barton and Etheridge, 2011). After a few generations, multiple blocks will have survived, and will thereafter propagate approximately independently, according to the branching process that we describe. Since hybridisation was relatively recent, blocks are still rare. Reported block lengths are up to 0.25cM (Iasi et al. (2024), Fig.1E), so the total selection coefficient of a block (including any discrete loci) can be up to, say, 0.25% and be of the same order as recombination. Thus, we need not worry about the complex later stage, in which introgressing blocks interact with each other.

So, we expect to see an overlay of sets of blocks, each set deriving from a different F1 individual. There could have been a single lucky hybridisation, where introgressing blocks were further aided by selection. At the opposite extreme, there might have been an initial mixing of around 2% of the population, followed by random drift. With an effective population size in the thousands, this would imply some tens of contributing F1 individuals.

Assuming no overlap in the sets of blocks, a reasonable assumption in the absence of strong selection, these scenarios could be distinguished by the distribution of block length, and by heterogeneity in the proportion of Neandertal ancestry as we scan along the genome. The distribution of block length is influenced by demographic change and by variation in the timing of gene flow (Iasi et al., 2021), as well as by the number of selected loci: with one selected locus per block, length decreases by recombination alone, whilst with multiple selected loci, or in the infinitesimal limit considered here, blocks will be longer, or will survive intact if *s > r* with larger blocks predominating if *s >* 3*r*. The pattern of heterogeneity may be most informative, but it is not clear what the null expectation is: even with no selection, there will be considerable random variation in introgression. Selection on discrete loci will simply cause corresponding peaks in introgression (the usual interpretation), but even if selection is uniformly spread over an infinity of loci, the branching process still leads to extreme variation (e.g. Fig. 1). Several lines of evidence suggest selection on alleles introgressing from Neandertal: the deficit of Neandertal ancestry in constrained regions, and on the X chromosome (Petr et al., 2019); the overall decrease through time (Iasi et al., 2024); and enrichment of certain classes of gene (Zeberg et al., 2024). Nevertheless, it is not at all clear that the pattern of heterogeneity across the genome is inconsistent with descent from a limited number of hybridisations.

How polygenic is selection on an introgressing block likely to be? We know that complex traits are typically highly polygenic, perhaps even being influenced by most of the expressed genome (Boyle et al., 2017). Similarly, purifying selection maintains a few percent of sites in species such as humans and Drosophila (Charlesworth, 2015). Thus, populations that have diverged under selection, or that have accumulated distinct mutation loads, may differ at least at tens of thousands of sites. We do not expect these differences to be fixed, but what matters is the number of differences between a single introgressing genome, and the consensus genotype of the recipient population. Under the infinitesimal model, this divergence is expected to be close to neutral, and so can be estimated from genome-wide divergence.

Such arguments suggest that there may be selected divergence at very many sites, even within small blocks (*<* 0.1cM, say). However, our stronger assumption that selective effect is proportional to block length will break down as blocks become small, and the idiosyncratic effects of the alleles they carry become important. We view our analysis as a tractable starting point for more realistic models of the relation between selection and block length. Moreover, we emphasise that key results – in particular, the increase of intact blocks when selection is stronger than recombination – are not sensitive to numbers of loci or interactions between them.

Our analysis was based on the strong assumption that introduced blocks have a selective advantage proportional to their map length, so that the ratio *s/r* is constant. In reality, even if alleles have additive effects, these effects will vary along the genome. Sachdeva and Barton (2018a) considered an infinitesimal limit, in which the net effects of very many alleles are Gaussian, and the key parameter is now the variance of effects per map length. Analysis of this case is difficult, because the effects of new recombinant blocks follow a distribution, rather than being determined by their length, as in our analysis. Moreover, blocks generated by independent crossovers will have correlated effects if they overlap; even simulation is complicated by the need to track these correlations. In this regime, the blocks that emerge are those with the maximum net rate of increase, *s* − *r*; these tend to be of intermediate length. The ratio *s/r* that has been of central importance in the case of linear selection that we have studied here, is no longer the key parameter when selection and recombination rates do not increase in precisely the same way with block length. Sachdeva and Barton (2018a) argue that in their setting, for an initial block of length *x*, one expects surviving blocks to contribute around 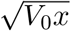 to the trait value, where *V*_0_ is the genic variance per unit map length. This is large enough to have a strong advantage, but not so large as to be broken up by recombination. Even in generalisations that include epistasis, it remains true that any block with net advantage stronger than recombination can increase intact, and the block with the highest net rate will dominate. Note that even two discrete alleles with selective values *s*_1_ and *s*_2_ will increase together if *s*_1_ + *s*_2_ − *r*_12_ *> s*_1_, *s*_2_ (where *r*_12_ is the recombination rate between them), so that it will be hard to distinguish two discrete loci from very many spread over the same region.

How efficient is selection in the presence of recombination? We analyse the initial invasion into a large population of a uniformly favourable block of genome. This would fix most rapidly if there were no recombination: the positive linkage disequilibrium between loci accelerates their increase. However, even when recombination is faster than selection (*r > s >* 0) a block of length *x* of introduced material has probability 2*sx* of being established at least in part, and its mass increases even as favourable blocks are broken up. Once these blocks become common, they compete with each other to fix: they are now in negative linkage disequilibrium, and so recombination is advantageous, since it can bring together multiple blocks. We do not analyse this in any detail (it is a hard problem). In the case of a single allele in a very large population (log(4*N*_*e*_*s*) 1), most time is spent in the branching phase, and the sweep to fixation is relatively fast (1*/s* rather than (1*/s*) log(4*N*_*e*_*s*)). In our setting, it is much less clear what will happen. If the surviving blocks are sufficiently dispersed along the genome (*r > s*), then they may reproduce essentially independently, meaning that the branching phase lasts much longer than in the case of a single allele. On the other hand, we expect to see an increase of new (initially rare) recombinants, which could also be treated as a branching process, albeit evolving in a heterogeneous background. We can certainly hope that our analysis gives some insight into what will ultimately fix. This is bounded above by the material that survives – which in principle could well be fixed if it could recombine onto the same genome. For the case *s > r >* 0, we can establish a lower bound, set by the largest block that survives, which will grow faster than any other, and must ultimately fix. There may be other blocks that have also survived intact, but which grow more slowly. We expect that, since these exist in very large numbers, they will recombine to also fix (though this is not guaranteed).

In a more general setting, where positive and negative material is intertwined, it is not obvious what selection strength is optimal. Strong selection will rapidly establish the block with fastest net rate of increase (*s* − *r*) but this may include ‘trapped’ deleterious material. Recombinants carrying just the beneficial material may increase more slowly, so that in the end, the largest block will still fix. Weaker selection would allow more time for recombination to act, so that the two disjunct beneficial blocks could recombine together, giving a stronger ultimate gain.

We have found a complete solution for the initial increase of a block of genome that is under linear selection, but much remains to be done. We see three open tasks. First, to know how to extend the branching process analysis to more general forms of selection (including the infinitesimal model defined by Sachdeva and Barton (2018a)). Second, to exploit these branching process analyses to understand the ultimate response to selection after a new genome is introduced into a homogeneous population. Third, to understand how blocks of genome evolve in a heterogeneous population, in which every genome has different effects. These are hard problems, but as our analysis here illustrates, one can sometimes make progress by combining a branching process with diffusion approximations.

## A Appendix

This appendix contains the derivations of the results presented in Section 3, as well as a justification of the diffusion approximation.

### A.1 Diffusion approximation

A solution to (2.1) can be formally defined by prescribing the dynamics of the integral of the measure-valued process (*Z*_*t*_)_*t*≥0_ against regular test functions. Let ⟨*Z*_*t*_, *ϕ⟩* := ∫ *ϕ*(*x*)*Z*_*t*_(d*x*) and define a function *L*[*ϕ*] as

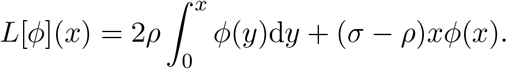

The integral operator *L* is the adjoint of the operator *L*^∗^ corresponding to the deterministic part of (2.1), in the sense that for a measure *µ*

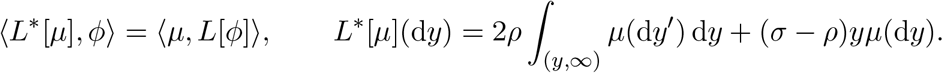

We say that (*Z*_*t*_)_*t*≥0_ is a solution to (2.1) if, for any *ϕ* regular enough,

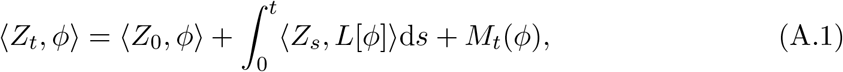

where *M*_*t*_(*ϕ*) is a continuous martingale with quadratic variation

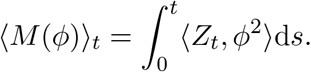

Intuitively, (A.1) indicates that the increments of ⟨*Z*_*t*_, *ϕ⟩* are the sum of two contributions: a deterministic part ⟨*Z*_*t*_, *L*[*ϕ*] ⟩ which combines the effect of recombination and selection; and a random fluctuation due to births and deaths with infinitesimal variance ⟨*Z*_*t*_, *ϕ*^2^⟩. We refer to Perkins (2002), Section II.5 for a rigorous treatment.

In the probabilistic literature, a solution to (2.1) is also called a measure-valued branching process or superprocess, see e.g. Etheridge (2000) for an introduction. These objects can be conveniently studied using Laplace transforms, which will be our main tool to derive the results presented in the main text. Under some regularity assumptions, the Laplace transform of (*Z*_*t*_)_*t*≥0_ can be expressed as

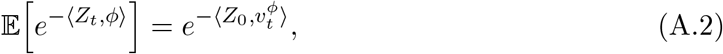

where, for a non-negative function *ϕ* on 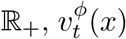 solves the non-linear evolution equation

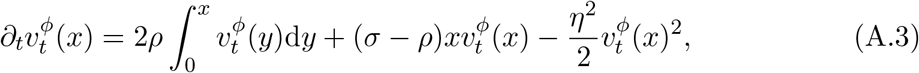

with initial condition 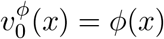. A rigorous proof of (weak) convergence of the sequence of scaled branching processes considered here to the measure-valued diffusion (*Z*_*t*_)_*t* ≥ 0_ would follow the standard pattern for construction of superprocesses, for full details of which we refer to Perkins (2002, Chapter II), or Etheridge (2000, Section I.1.5). Here we satisfy ourselves with computing the infinitesimal mean and variance of our scaled branching process, and showing that they converge to those implicit in (2.1).

Writing *Y*_*t*_(d*y*) for the number of blocks of the branching process with lengths in the infinitesimal interval [*y, y* + d*y*) in generation *t*, and recalling the definition of the scaled parameters *r* = *ρ/K, s* = *σ/K*,

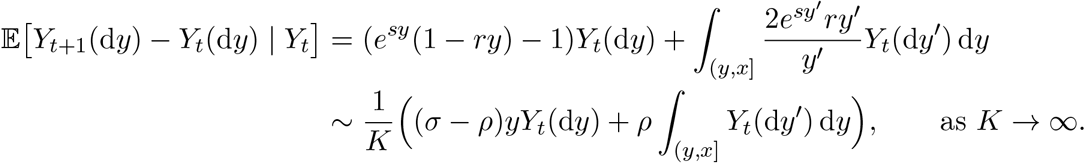

The first term is the increment due to the average number of copies of itself that a block produces, and the second term accounts for the creation of blocks with length in [*y, y* +d*y*) through recombination of larger blocks. Similarly, the infinitesimal variance is computed as

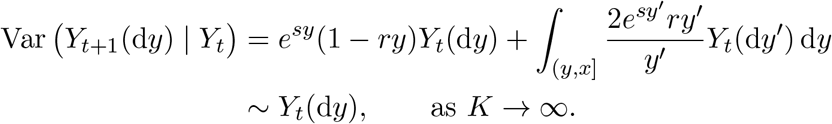

Since the number of offspring is distributed as a Poisson point process, if *y* ≠ *y*′

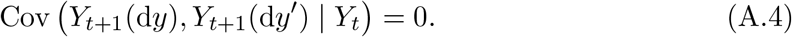

Therefore, if we introduce the rescaled process 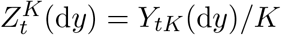, the computations above show that, as *K* → ∞,

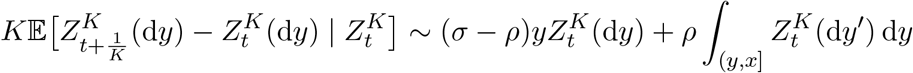

and

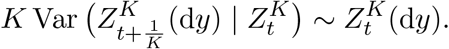

In turn this suggests that, if 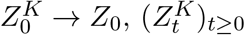 converges to the solution of

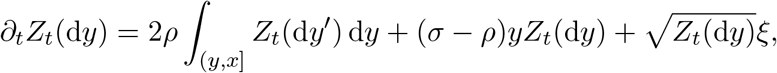

where the fact that *ξ* is a space-time white noise comes from the absence of correlation noted in (A.4). Taking the initial condition *Z*_0_ = *δ*_*x*_, a Dirac delta function at *x*, corresponds to starting our branching process from *K* individuals at time zero, each of which carries a block of length *x*, but assigning a ‘mass’ 1*/K* to each individual.

Finally, let us outline how this calculation should be adapted to derive (2.1) when the offspring number is no longer Poisson distributed. We only illustrate the computation of the (co-)variance of block lengths after the first generation. Recall that we denoted by *P* the number of ‘parental pairs’ in which an individual is involved. Since the probability that a recombination event occurs within the block is small, up to an error term *O*(1*/K*) we can assume that each of the *P* offspring inherits the parental block with probability 1*/*2. Thus, for any test function *ϕ* conditioning on *P* shows that

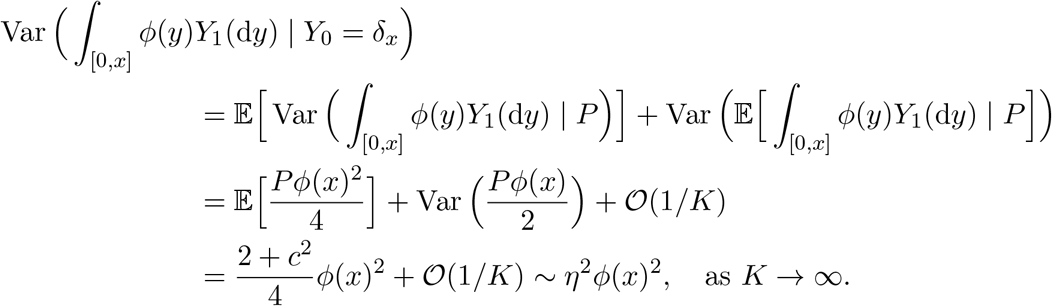

By the Cauchy–Schwartz inequality and choosing *ϕ*(*z*) = **1**{*z* ∈ [*y, y* + d*y*)}, for *y* ≠ *y*′,

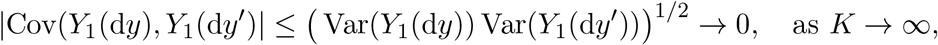

so that the noise terms for different block lengths are uncorrelated in the limit.

### A.2 Survival probability

We will use the diffusion approximation *Z*_*t*_ started from *δ*_*x*_*/K* to write down an approximation for *Y*_*tK*_. The limit (*Z*_*t*_)_*t*≥0_ enjoys the branching property, meaning that if 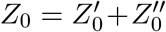 the process (*Z*_*t*_)_*t*≥0_ is distributed as the sum of two independent copies of itself, started from 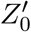 and 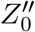. The branching property implies that

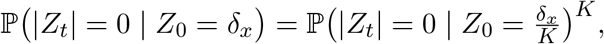

where |*Z*_*t*_| is the total mass of the diffusion at time *t*. Using this identity,

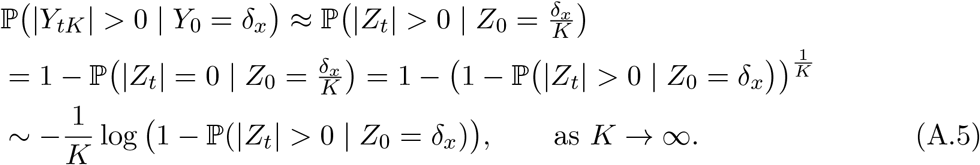

Let us introduce (with a slight abuse of notation relative to the main text)

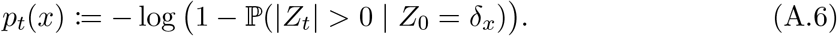

To approximate the survival probability of our branching process, we study the limit of *p*_*t*_(*x*).

Before proceeding with establishing the asymptotic behaviour of *p*_*t*_(*x*), let us illustrate how our approximation will work in an example. This will provide some justification for our abuse of notation. We shall see below that if selection is negative (*s <* 0, and therefore *σ <* 0), then

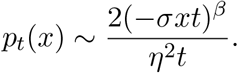

Our strategy above for approximating our branching process at large times says that to estimate ℙ (|*Y*_*t*_ | *>* 0), we should substitute *t/K, ρ* = *rK*, and *σ* = *sK* in this expression and multiply by 1*/K* to obtain

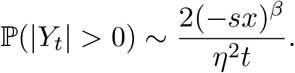

In particular, our large parameter *K* has cancelled. This will be a feature of all the approximations for our branching process that we obtain from the diffusion approximation, but it is important to remember that, in order for them to be valid, we are assuming that *r* and *s* are small, and *t* is large enough that if *r >* 0 (resp. *s >* 0) *rt* » 1 (resp. *st* » 1).

We estimate *p*_*t*_(*x*) as defined in (A.6) from the evolution equation. Let *v*^*θ*^(*x*) be the solution to (A.3) with initial condition *ϕ* ≡ *θ*. Observe that since (*Z*_*t*_, *θ*) is monotone in *θ*, 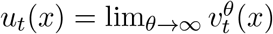 exists and

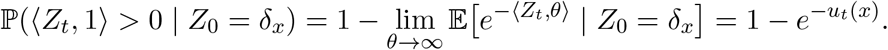

Comparing to (A.6), we see that *p*_*t*_(*x*) = *u*_*t*_(*x*). The limit (*p*_*t*_(*x*)) will still solve the evolution equation (A.3),

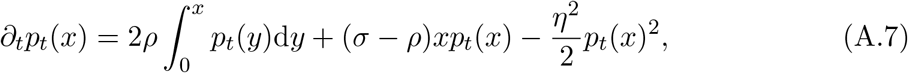

but started ‘from infinity’ at time 0.

To estimate the asymptotic behaviour of the survival probability, we shall assume that *p*_*t*_(*x*) varies slowly with respect to time. Setting ∂_*t*_*p*_*t*_(*x*) = 0 in (A.7) and taking a derivative with respect to *x*, we see that in this case *p*_*t*_(*x*) solves (approximately) the differential equation

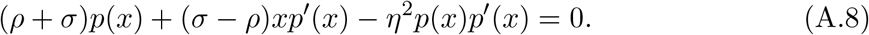

Moreover, setting *x* = 0 in (A.7),

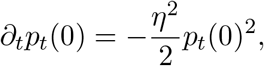

whose unique solution satisfying *p*_0_(0) = 1 and *p*_*t*_(0) → ∞ as *t* ↓ 0 is *p*_*t*_(0) = 2*/*(*η*^2^*t*). Of course, this corresponds to the survival probability up to time *t* of a single neutral locus. We approximate *p*_*t*_(*x*) by the solution to (A.8) with initial condition *p*_*t*_(0) = 2*/*(*η*^2^*t*).

Equation (A.8) can be solved by transforming it into an exact differential equation using an integrating factor *I*(*p*). More precisely, if we can find *I*(*p*) such that

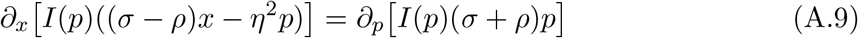

then the differential

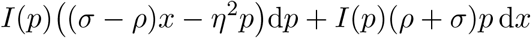

on (0, ∞) × (0, ∞) is closed. By Poincaré’s Lemma, see for instance Weintraub (1997, Chapter VI, Lemma 1.14), it is an exact differential, which means that there exists *F* such that

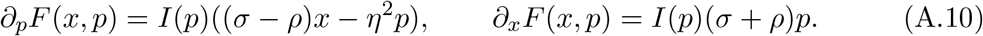

Observe that for any solution *p* of (A.8)

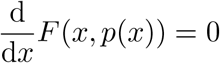

and therefore *F* (*x, p*(*x*)) = *c* for some *c* ∈ ℝ. Equation (A.9) can be written as

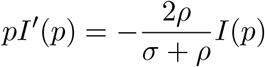

and we choose as integrating factor 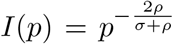. Integrating the differential d*F* along two straight lines, one going from (0, *p*(0)) to (0, *p*(*x*)), the other from (0, *p*(*x*)) to (*x, p*(*x*)),

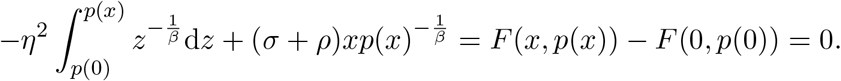

When *σ* ≠ 0 and so *β* ≠ 1, this equation becomes

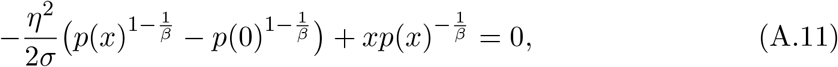

and when *σ* = 0, so *β* = 1,

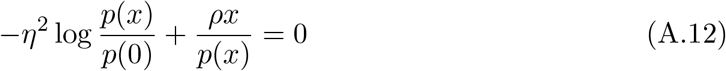

Substituting *p*(0) = 2*/*(*η*^2^*t*) in this equation provides an implicit expression for *p*(*x*). We consider the cases *σ* = 0, *σ <* 0, and *σ >* 0 separately.

#### Neutrality (*σ* = 0)

We quickly show how to recover the results in Baird et al. (2003) for the neutral case *σ* = 0. Introducing *q*_*t*_(*x*) such that 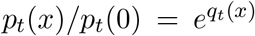, equation (A.12) becomes

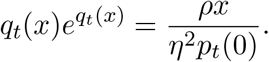

The solution to this equation can be expressed using Lambert’s *W* function, which is implicitly defined as the solution to *W* (*z*)*e*^*W* (*z*)^ = *z*. Namely,

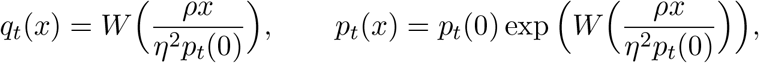

Using that *W* (*z*) = log *z* − log 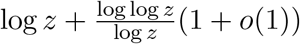 as *z* → ∞, we obtain that

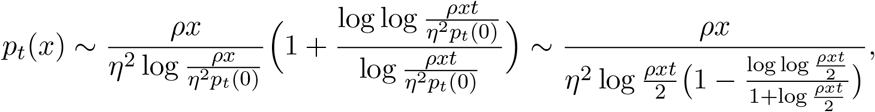

which is equation (7) in Baird et al. (2003). Substituting *ρ* = *Kr*, replacing *t* by *t/K* and multiplying the whole expression by 1*/K* to recover the survival probability for our branching process, we find

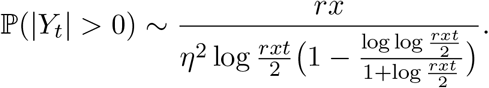

#### Negative selection (*σ <* 0)

For *σ <* 0, a comparison with the neutral case shows that *p*_t_ (*x*) → 0 as *t* → ∞. Hence,^1^ neglecting the lower order term 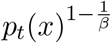 in (A.11), simple manipulations show that

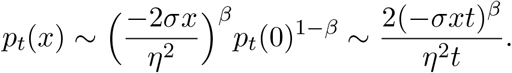

For the branching process we recover,

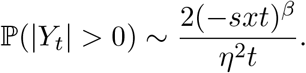

#### Positive selection (*σ >* 0)

Lastly, since *σ >* 0 implies that 1*/β <* 1, the term 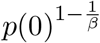 in (A.11) is negligible for large *t*. The probability *p*_*t*_(*x*) converges to the solution of

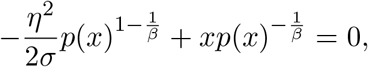

which is *p*^∗^(*x*) = 2*σx/η*^2^. We can compute the second order term in the asymptotic expansion of the survival probability by introducing *q*_*t*_(*x*) = *p*_*t*_(*x*) − *p*^∗^(*x*). Since *q*_*t*_(*x*) « 1, performing a Taylor expansion in (A.11) yields that

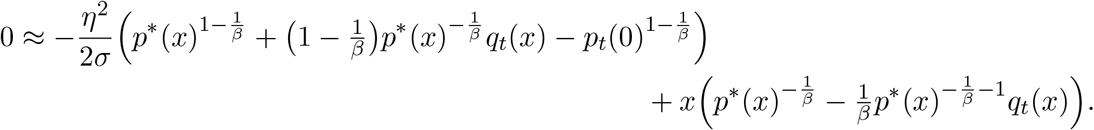

Rearranging,

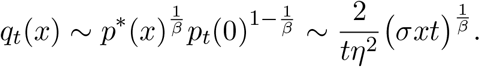

Adding *p*^∗^(*x*), substituting *σ* = *Ks, ρ* = *Kr*, replacing *t* by *t/K* and multiplying the whole expression by 1*/K* to recover the survival probability for our branching process, we find

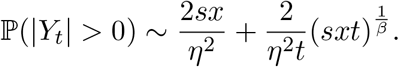

As remarked in the main text, at first sight this result is surprising. To see that it makes sense mathematically, one can use equation (A.1) to see that d ⟨*Z*_*t*_, *x* ⟩ = *s* ⟨*Z*_*t*_, *x*^2^⟩ d*t* + d*M*_*t*_, where the infinitesimal variance of the fluctuation *M*_*t*_ is ⟨*Z*_*t*_, *x*^2^⟩ d*t*. Since this is (up to the constant *s*) the same as the infinitesimal change in the drift, the total length of blocks is a random time change of drifted Brownian motion, and the probability that the drifted Brownian motion never hits zero started from the point *x* is exp(−2*sx*). This isn’t enough to prove the result, as we would also need to show that as *t* → ∞ the corresponding time change *τ* (*t*) → ∞, but at least mathematically it demystifies it a little.

### A.3 Survival probability of a distinguished locus

We turn to the calculation of the probability of survival of a distinguished locus within the introgressing block. Write *p*_*t*_(*x*_1_, *x*_2_) for the probability that a locus that initially has length *x*_1_ of introgressing genome to its left, and length *x*_2_ to its right, survives until time *t*. Exactly as for the case in which we do not distinguish a particular locus, using the diffusion approximation we find that *p*_*t*_(*x*_1_, *x*_2_) can be approximated by the solution to

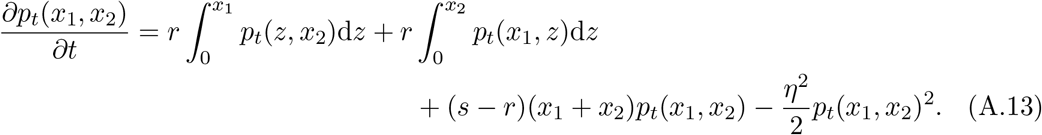

Finding an approximation to the solution to this equation is significantly more challenging than the previous case. However, we can use the method of the previous section to estimate *p*_*t*_(0, *x*), the probability that the left endpoint of the introgressing block survives up to time *t*.

To ease notation, we write *p* (*x*) = *p*_t_ (0, *x*). Equation (A.7) is replaced by

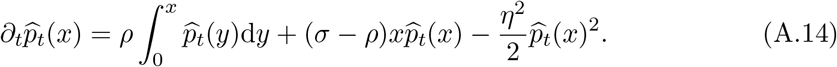

As before, our strategy is to assume that 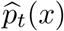 varies slowly with respect to time, set 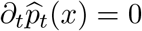 and differentiate with respect to *x*. Equation (A.8) is replaced by

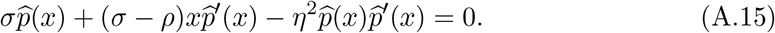

We look for the solution 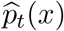 to (A.15) with initial condition *p*_*t*_(0) = *u*_*t*_(0) = 2*/*(*η*^2^*t*).

The integrating factor, 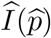 that turns equation (A.15) into an exact differential equation satisfies

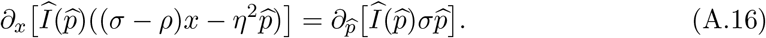

Solving, we find 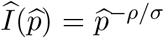. Once again Poincaré’s Lemma guarantees the existence of 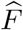 such that

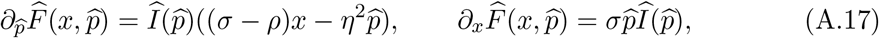

and for any solution 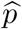 of (A.15)

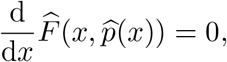

and therefore 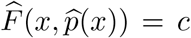 for some *c* ∈ ℝ. Integrating the differential 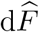 along two straight lines, one going from 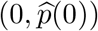 to 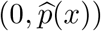, the other from 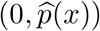 to 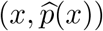,

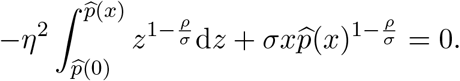

That is

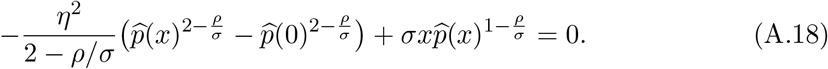

If *σ > ρ/*2, then 2 − *ρ/σ >* 0 and so when we substitute 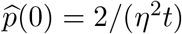, the term involving 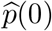 will become negligible as *t* → ∞. Rearranging the other terms we obtain

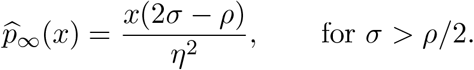

Reverting to the original units for our branching process, if 2*s* − *r >* 0, then the probability of survival of the left endpoint is approximately (2*s* − *r*)*x/η*^2^.

On the other hand, if *σ < ρ/*2, the blocks containing the left endpoint locus will eventually die out and so ignoring the term involving 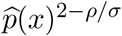 and rearranging, we arrive at

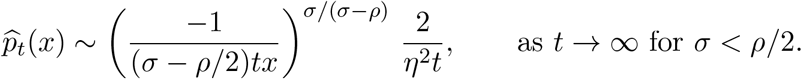

Reverting to the units of our branching process, we find the probability that the left endpoint of the introgressing block survives until time *t* when 2*s* − *r <* 0 is

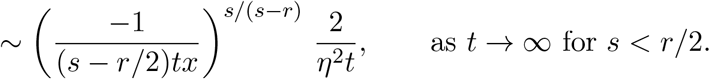

We have been unable to find a good approximation for *p*_*t*_(*x*_1_, *x*_2_) when both *x*_1_ and *x*_2_ are nonzero. As explained in the main text, recombination is less effective at shedding block length linked to the distinguished locus if that locus is at an interior point and so, for example, in the case 2*s* − *r >* 0 one expects that *p*_∞_ (*x*_1_, *x*_2_) − *p*_∞_ (0, *x*_1_ + *x*_2_) *>* 0. Simulations show that the difference is not huge, and indeed by comparison with the case *r* = 0, we see immediately that *p*_∞_ (*x*_1_, *x*_2_) ≤ 2*s*(*x*_1_ + *x*_2_)*/η*^2^. We now establish a lower bound on *ε*(*x*_1_, *x*_2_) := *p*_∞_ (*x*_1_, *x*_2_) − *p*_∞_ (0, *x*_1_ + *x*_2_). Write *p*_0_(*x*_1_, *x*_2_) = (2*s* − *r*)(*x*_1_ + *x*_2_)*/η*^2^. Then a straightforward calculation shows that

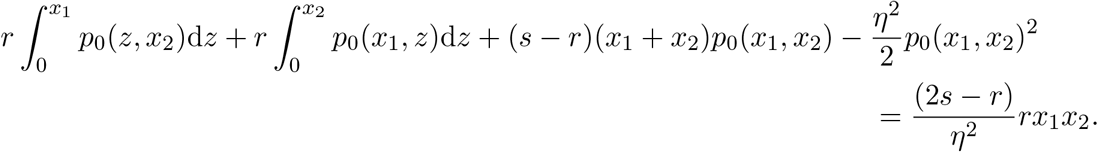

On the other hand substituting *p*^∞^ (*x*_1_, *x*_2_) into the same equation (which is equation (A.13) with the left hand side set to zero) gives zero, and so taking the difference,

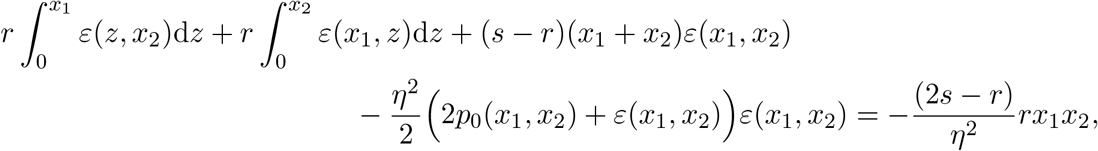

where we have used that *p*^∞^ (*x*_1_, *x*_2_) = *p*_0_(*x*_1_, *x*_2_) + *ε*(*x*_1_, *x*_2_) in the last term on the left hand side. Rearranging,

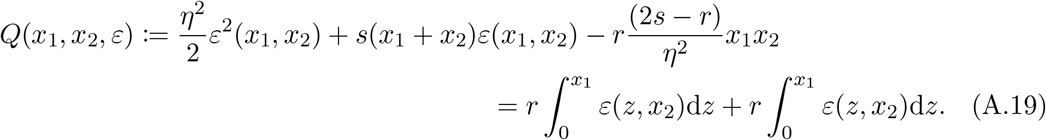

The quadratic on the left is negative when *ε* = 0, and the expression on the right is positive, so we can bound *ε*(*x*_1_, *x*_2_) below by the positive root of *Q*(*x*_1_, *x*_2_, *ε*) = 0. That is

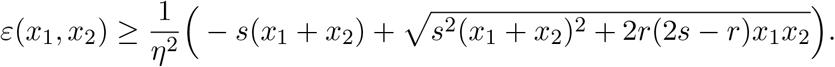

Expanding the square root, if *r/s* is small,

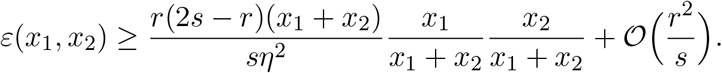

One could do a more refined analysis by substituting this lower bound in the right hand side of (A.19) to obtain a new quadratic and iterating, but this does not lead to an elegant expression.

### A.4 Length of the largest block

In this section, we assume that *s > r*, so that blocks of macroscopic length (that is of length of the same order as that of the initial block) can survive. We estimate the distribution of the largest block that survives from the diffusion approximation.

Let us denote by 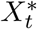 the length of the largest block at time *t* in the diffusion approximation (*Z*_*t*_)_*t*≥0_. Formally, *Z*_*t*_ is a random measure on (0, ∞) and the largest block is the supremum of its support. Namely, 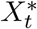 is the unique random variable such that

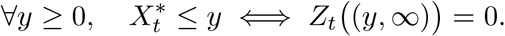

Note that this identity only holds provided that we let 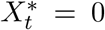 if *Z*_*t*_ = 0, which is a convention that we enforce from now on. In other words, the length of the largest block is arbitrarily set to 0 on the event that the population is extinct.

Let us write 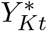 for the longest surviving block in the branching process at time *Kt*. Our strategy for approximating its distribution from that of the longest block in the diffusion approximation mimics our approach in Section A.2. Using the branching property,

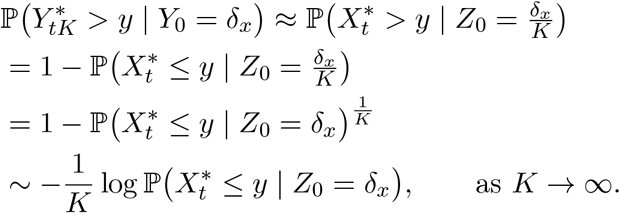

We therefore study the limit as *t* → ∞ of

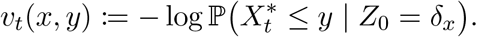

If *v*_*t*_(*x*; *ϕ*) denotes the solution to the evolution equation (A.3) started from *v*_0_(*x*; *ϕ*) = *ϕ*(*x*) and we set *ϕ*_*y*_(*z*) = **1**_{*z>y*}_, then

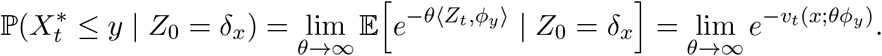

We are going to abuse notation and define

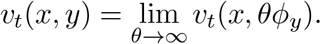

For each fixed *y*, the function *v*_*t*_(*x, y*) is again a solution to the evolution equation (A.3), but started from the initial condition

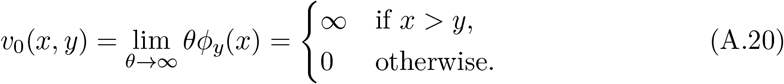

Letting *t* → ∞, we deduce that

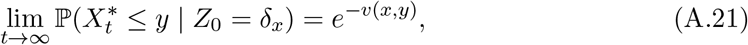

where *v*(*x, y*) = lim_*t*→∞_ *v*_*t*_(*x, y*) is a stationary solution to (A.3), that is,

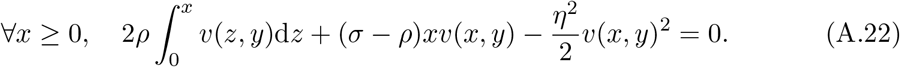

Equation (A.22) does not have a unique solution and we need to specify an appropriate boundary condition. By (A.20), we know that *v*_*t*_(*x, y*) = 0 if *x* ≤ *y*. Moreover,

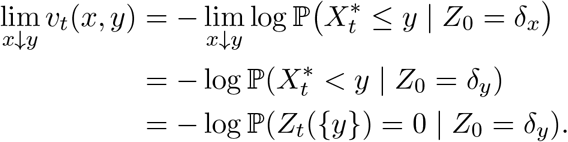

The quantity 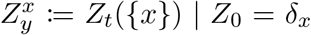 is just the diffusion approximation for the number of copies of the block *x* that are passed on intact, and so it behaves as a single locus with fitness (*σ* − *ρ*), for which

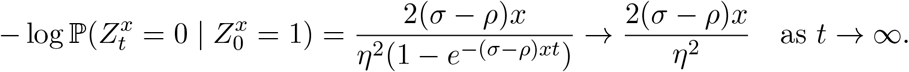

In particular, we see that *v*_*t*_(*x, y*) is discontinuous at *x* = *y*. Letting *t* → ∞, we look for a solution to (A.22) for which *v*(*x, y*) = 0 for *x* ≤ *y* and

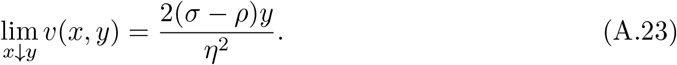

We have already established in Section A.2 that solutions to (A.22) can be implicitly represented as *F* (*x, v*(*x, y*)) = *c*_*y*_, for some *c*_*y*_ ∈ ℝ and where the derivatives of *F* are prescribed in (A.10). Therefore, for *y < x*′ *< x*, by integrating the differential of *F* along two straight lines, one going from (*x, v*(*x*′, *y*)) to (*x, v*(*x, y*)), and the second from (*x*′, *v*(*x*′, *y*)) to (*x, v*(*x*′, *y*)), we obtain that

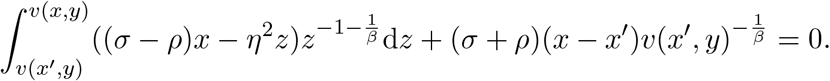

Letting *x*′ ↓ *y* and using (A.23),

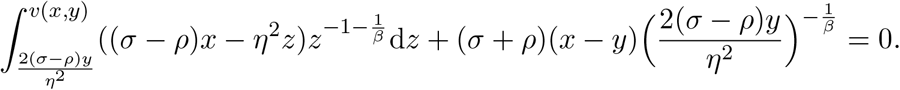

The integral can be simplified further by using that an antiderivative of the integrand is

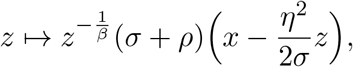

from which a direct computation shows that *v*(*x, y*) satisfies

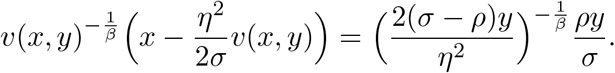

We seek 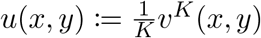, where *v*^*K*^(*x, y*) is obtained by substituting *σ* = *Ks, ρ* = *Kr*. By introducing

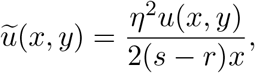

this equation simplifies to

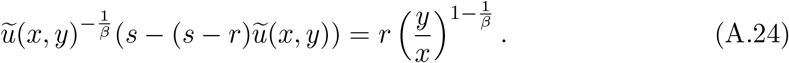

Noting that 0 *<* −1*/β <* 1, define the map 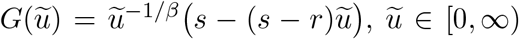. It satisfies 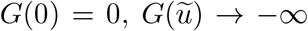 as 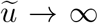, and, differentiating, it has a unique critical point (*G*′(1*/*2) = 0), so equation (A.24) always has two solutions. Moreover, since *y* ≤ *x*, and *G*(1) = *r*, one of those solutions will be less than 1*/*2 and the other at least 1. Evidently *u*(*x, y*) is monotone decreasing in *y*, and as *y* ↑ *x, u*(*x, y*) approaches the probability that a block of length *x* leaves intact copies of itself in the indefinite future, while *u*(*x*, 0) is the probability that it leaves any genetic descendants at all, so

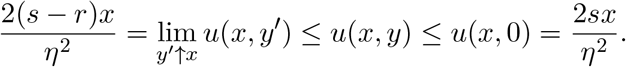

Therefore, 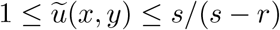. We conclude that

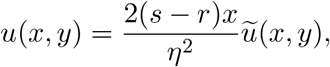

where 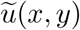 is the (unique) solution of (A.24) taking values in [1, ∞).

### A.5 Expected density of blocks

We now turn to the expected density of blocks *m*_t_ (*x*, d*y*). Recall that 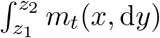) gives the expected number of blocks with lengths in the interval [*z*_1_, *z*_2_] that are present in the population at time *t*, given that we started from a single block of length *x* at time zero. Once again we use the diffusion approximation, and exploit the branching property:

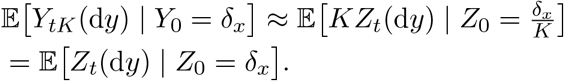

Therefore, the mean measure of the superprocess at time *t* provides an approximation of the mean block length distribution at generation *Kt* and to recover expressions for the asymptotic behaviour of the mean measure of the branching process, we simply replace all occurrences of *σt, ρt* in the asymptotic expressions for the diffusion approximation by *st, rt*, respectively. Just as when we calculated survival probabilities, we shall abuse notation, this time defining

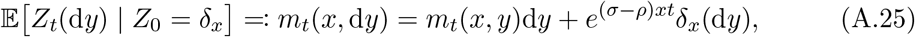

with the final form of the expressions that we obtain providing some justification.

Formally taking taking expectations on both sides of (2.1), the density *m*_*t*_(*x, y*) of the mean measure of the superprocess on (0, *x*) solves

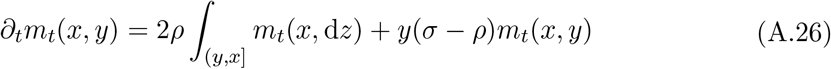

started from *m*_0_(*x, y*) = 0 (since *m*_0_(*x*, d*y*) = *δ*_*x*_(d*y*)), and note that *m*_*t*_(*x, x*) = *e*^(*σ*−*ρ*)*xt*^*δ*_*x*_(d*y*). Let us introduce the notation *M*_*t*_(*x, y*) = ∫ _[*y,x*]_ *m*_*t*_(*x*, d*z*) for the mean number of blocks with length larger than *y* at time *t*. Clearly, *M*_0_(*x, y*) ≡ 1 and, integrating by parts,

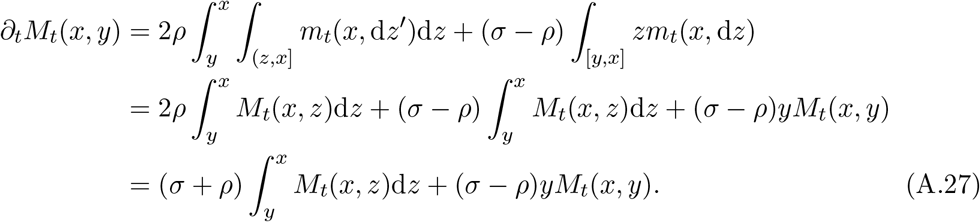

Define the Laplace transform of *M*_*t*_(*x, y*) with respect to time as

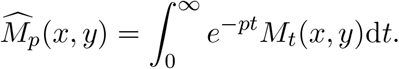

From (A.27), it solves

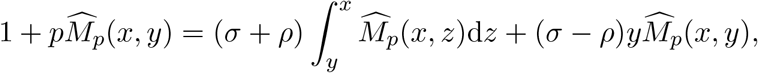

and differentiating both sides with respect to *y*,

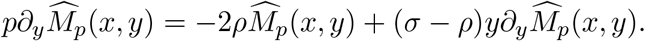

Rearranging the terms,

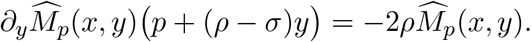

Moreover, since *M*_*t*_(*x, x*) = *e*^(*σ*−*ρ*)*xt*^,

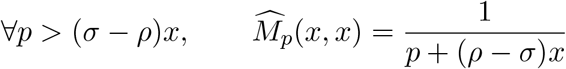

and therefore

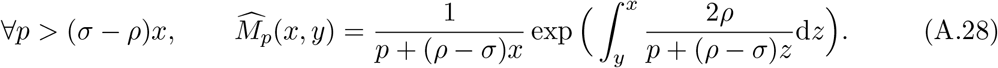

When *σ* ≠ *ρ*, (A.28) becomes

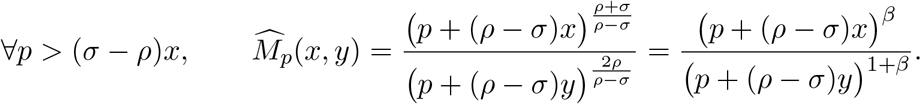

This Laplace transform can be inverted explicitly in terms of the confluent hypergeometric function

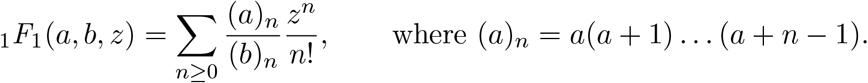

This relies on the elementary identity that

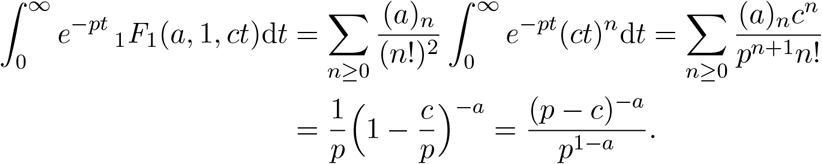

Choosing *a* = 1 + *β* and *c* = (*ρ* − *σ*)(*x* − *y*),

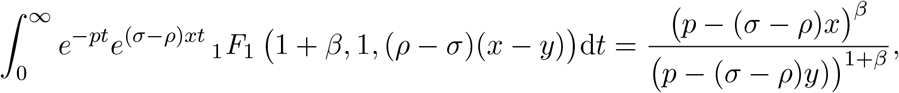

from which we deduce that

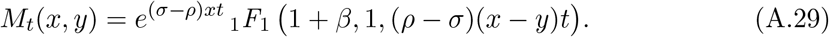

Differentiating with respect to *y* yields the corresponding expression for the density

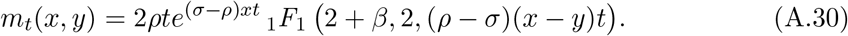

Substituting *ρ* = *Kr, σ* = *Ks*, and evaluating this expression at *t/K* to recover approximate expressions for the branching process gives (3.6) and (3.10) in the main text.

The known asymptotic behaviour of _1_*F*_1_(*a, b, z*),

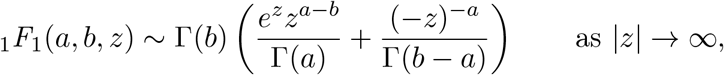

now provides expressions for the asymptotic behaviour of *M*_*t*_(*x, y*), depending on whether we are in the sub- or supercritical regime.

#### Case *σ > ρ*

When *σ > ρ*, we deduce from

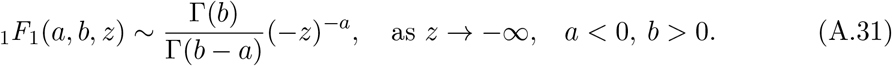

that

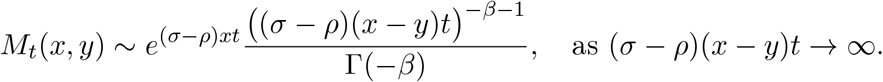

Setting *y* = 0 and substituting *st, rt* for *σt, ρt*, gives (3.8), and (3.9) follows readily.

#### Case *σ < ρ*

When *σ < ρ*, using that

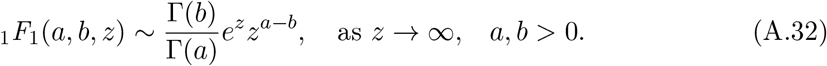

gives

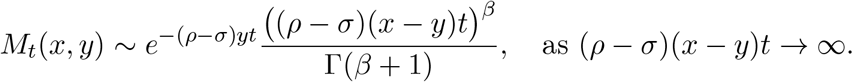

Setting *y* = 0 (and substituting *st, rt* as before) yields (3.13), and if *y* « *x*,

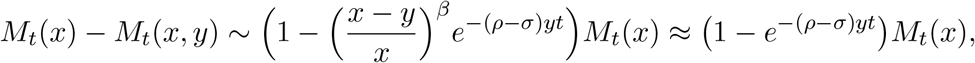

giving (again on substituting *st, rt* for *σt, ρt*) (3.14).

#### Case *σ* = *ρ*

When *σ* = *ρ*, (A.28) becomes

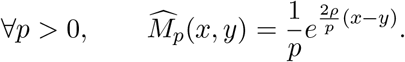

Recall the Laplace transform of a modified Bessel function of the first kind,

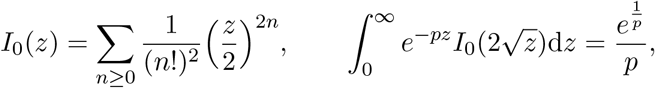

from which we can read off that

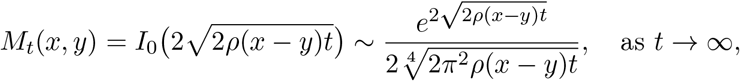

where we have used that

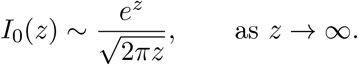

Setting *y* = 0 gives (3.17). For *y* « *x* and *t* » 1, (3.18) follows from

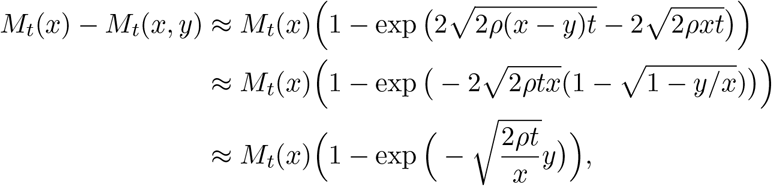

where as usual to obtain the corresponding expressions for our branching process we simply replace *ρt* by *rt*.

Finally, let us compute the average total mass. Integrating by part

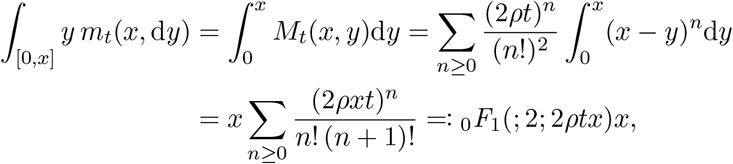

where _0_*F*_1_ is a confluent hypergeometric limit function. Again, substituting *rt* for *ρt* gives

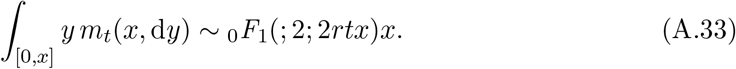

### A.6 Coalescence times

We now use our expressions for the expected number of blocks alive at a future time to derive an approximation for *C*_*t*_(*x, κ*), the expected number of pairs of individuals in generation *t* for which their most recent common ancestor lived before generation *κ*, given that the length of the original introduced block was *x*. Since we are going to use the expressions that we already obtained using the diffusion approximation for expected numbers of blocks in a given generation, we are implicitly assuming that *r* and *s* are small, but *t* is large (*rt* » 1 and, if *s* ≠ 0, *st* » 1). However, we work in natural (branching process) units (*s, r, t*). Under these conditions, we can, and do, approximate summation over discrete generations by integration with respect to time.

The most recent common ancestor of a pair of blocks at time *t* lived at time *τ* if the two blocks of this pair descend from two distinct offspring of the same individual living at *τ*. An individual carrying a block of length *y* at time *τ* produces a random number of copies of itself with mean 1 + *O*(1*/K*) and variance *η*^2^ + *O*(1*/K*). Therefore, the mean number of pairs of distinct offspring that it produces is *η*^2^*/*2 (up to an error of order 1*/K*), and each offspring (independently, because of the branching property) leaves an expected number *M*_*t*−*τ*_ (*y*) of descendants in generation *t*.

Integrating over *τ* ≤ *κ* and *y* ∈ (0, *x*], we find

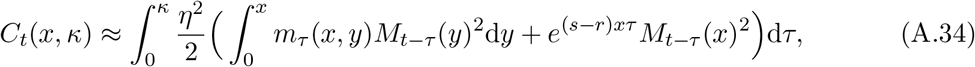

where *m*_*τ*_ (*x, y*) and *M*_*t*−*τ*_ (*y*) can be read off from (3.6) and (3.10) respectively for *s* ≠ *r*, and (3.17) and its derivative for *s* = *r*. Notice the second term in the integrand, corresponding to individuals carrying the whole of the original block.

Let us write

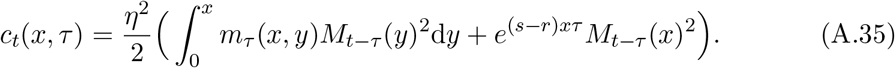

This is the quantity that we compute.

#### Weakly positive selection (*r > s >* 0)

First, when *r > s >* 0, using the estimate (3.13) for *M*_*t*−*τ*_ (*y*) and that (*t* − *τ*)^*β*^ ∼ *t*^*β*^, letting *t* → ∞ in (A.35) shows that

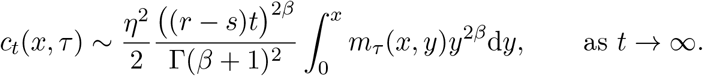

Replacing *m*_*τ*_ (*x, y*) by (3.10) and using the change of variable *z* = (*r* − *s*)(*x* − *y*)*τ*,

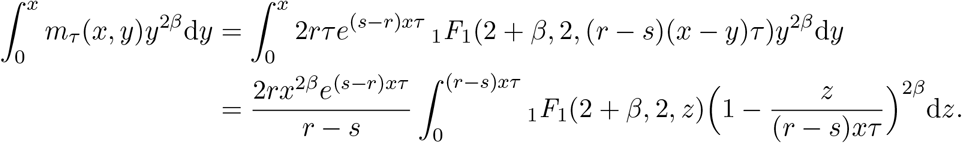

Therefore, introducing

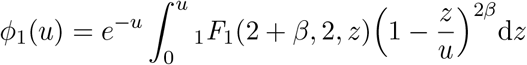

we obtain that

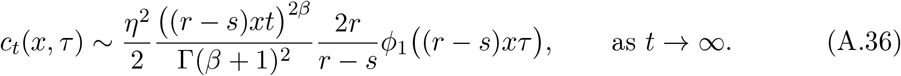

Now, note that *ϕ*_1_(*u*) → 0 as *u* → 0 and that, by (A.32),

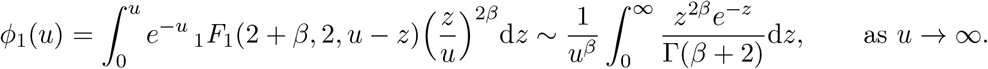

Since *β >* 1 (because 0 *< s < r*), the function

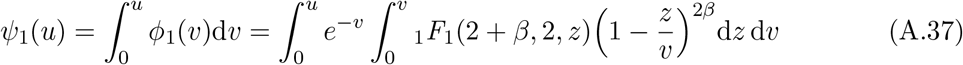

has a finite limit as *u* → ∞ and

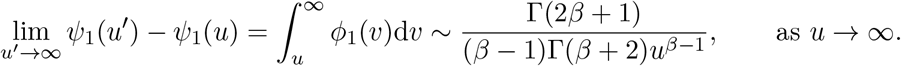

Taking integrals on both sides of (A.36),

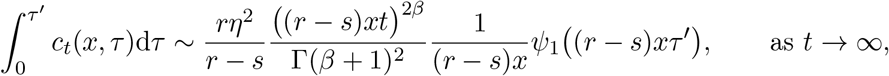

and so using (A.34)

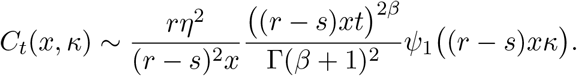

#### Negative selection (*s <* 0)

Again using the explicit expressions in (3.6) and (3.10),

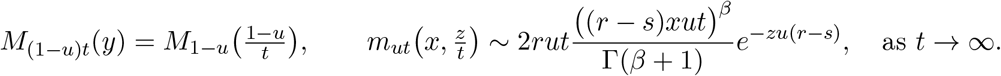

Substituting these in the expression for *c*_*t*_(*x, τ*) in (A.34), for any *u* ∈ (0, 1)

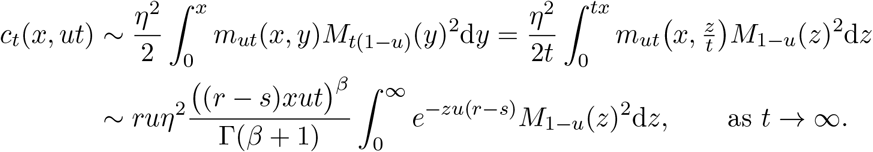

We now replace *M*_1−*u*_(*z*) by the expression in (3.6),

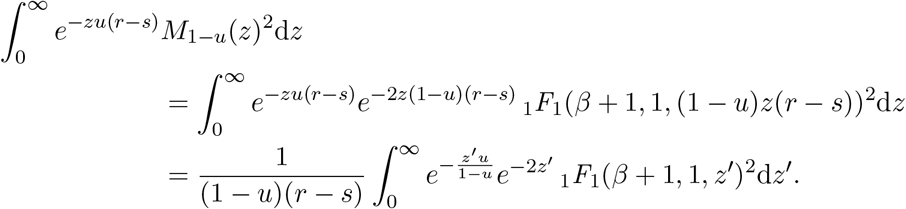

These two steps show that, for *s <* 0,

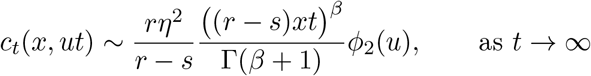

with

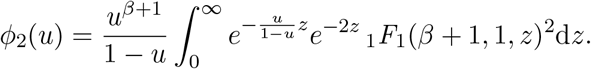

Now, (A.32) implies that *e*^−2*z*^ _1_*F*_1_(*β* + 1, 1, ∼ *z*)^2^ *z*^2*β*^*/*Γ(*β* + 1)^2^ as *z*→ ∞, from which we deduce that

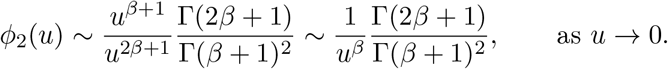

Since *β <* 1 (because *s <* 0), we can define

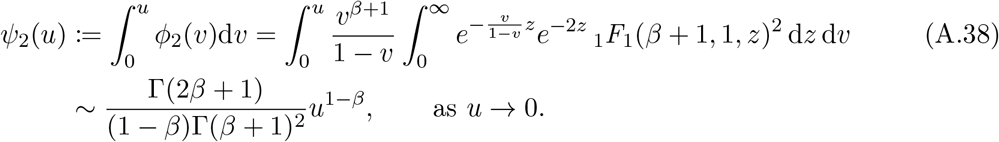

Altogether, we obtain that

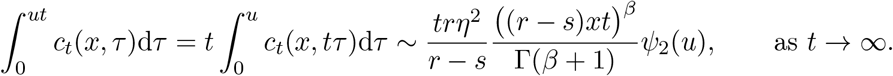

Finally, for *κ* = *ut*,

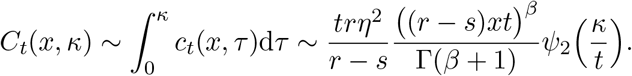

#### Neutrality (*s* = 0)

Fix *u* ∈ (0, 1). We see directly from equations (3.13) and (3.14) that, as *t* → ∞,

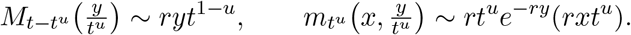

Substituting these in (A.35) and using the change of variable *y*′ = *yt*^*u*^

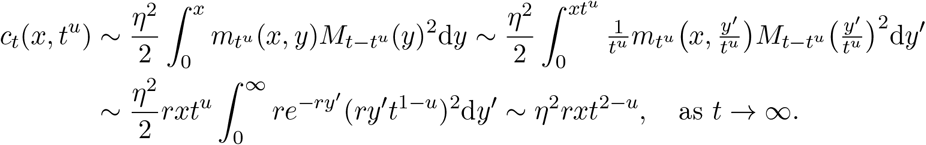

Therefore, making a change of variable *τ* = *t*^*u*^,

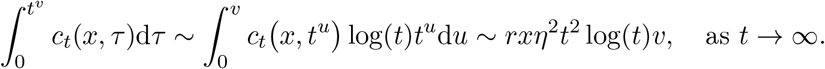

By (A.34), for *v* ∈ (0, 1),

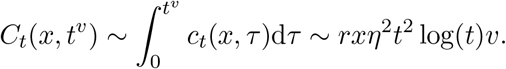

Thus if *κ* = *t*^*v*^,

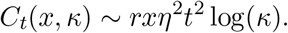

#### Strongly positive selection (*s > r*)

Finally, if *s > r*, we show that the leading order term in (A.35) is the second term. Using (3.8),

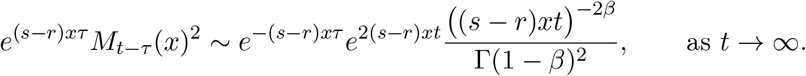

We show that the first term in (A.35) is negligible compared to *e*^2(*s*−*r*)*xt*^*t*^−2*β*^. Using (3.8) again,

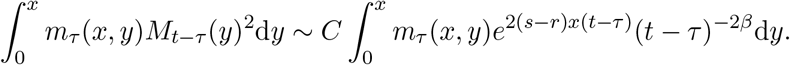

for some constant *C >* 0. Moreover,

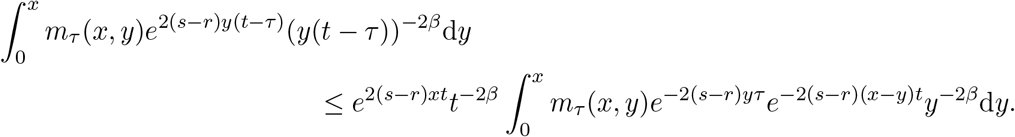

The above integral vanishes as *t* → ∞ by monotone convergence. Therefore,

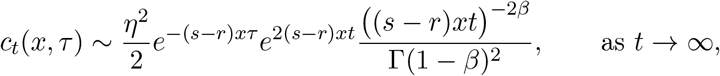

and

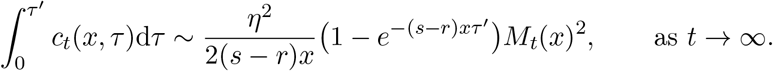

Finally,

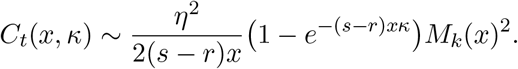

### A.7 Beneficial allele in a deleterious background

In this section we turn to the model with a major effect allele described in Section 2.4, and prove the results of Section 3.8.

#### Blocks with the beneficial allele

Recall that the diffusion approximation is the solution to (2.2) and let

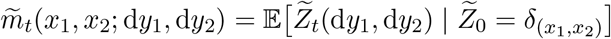

be the mean distribution of blocks in the diffusion approximation. It has a density on [0, *x*_1_) × [0, *x*_2_) that we also denote by 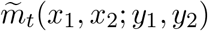. We also use the notation

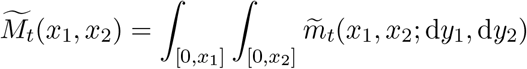

for the total number of blocks that contain the beneficial allele. Let us ease the notation and simpy write 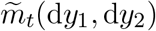. Taking expectations on both sides of (2.2),

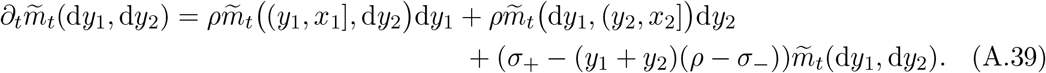

Now note that if 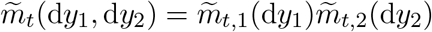 is a product measure, then

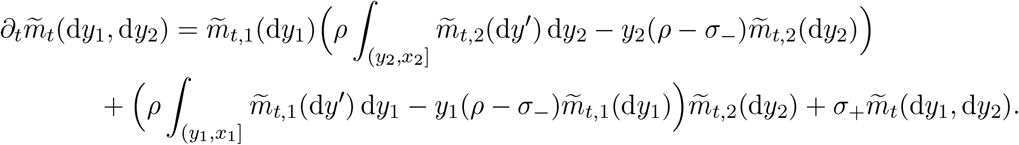

Since the initial block length distribution is a product measure 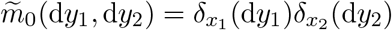, this computation indicates that 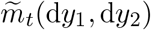 can be written as

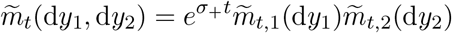

with

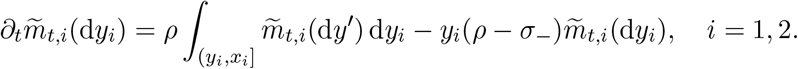

We see that this equation is of the same form as equation (A.26) for the density of blocks without the beneficial allele, up to replacing *ρ, σ*, and *β* in (A.26) by

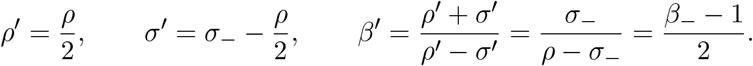

Therefore, by (A.29) the total mass is

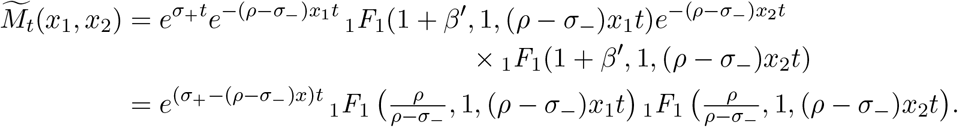

Similarly, by (A.30) the density has an exact expression,

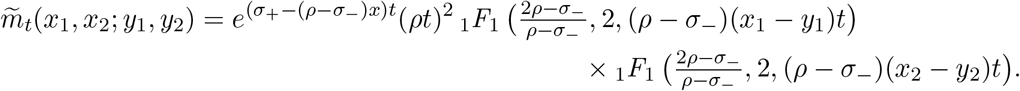

Letting *t* → ∞ using (A.32),

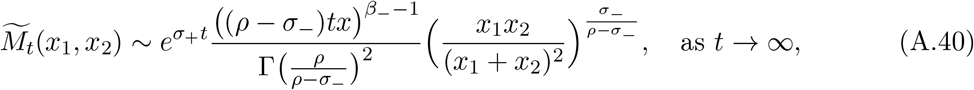

and

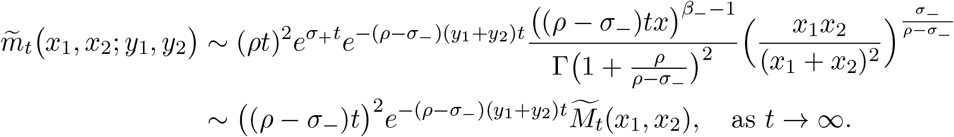

Integrating both sides yields

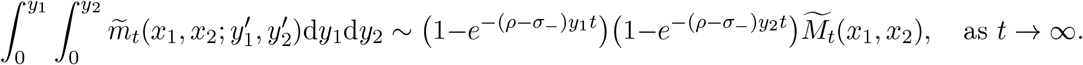

Rewriting these expressions in our original parameters *r* = *ρ/K, s*_−_ = *σ*_−_ */K, s*_+_ = *σ*_+_*/K*, at time *t/K* gives the results in the main text.

#### Blocks without the beneficial allele

We finally consider the dynamics of the purely deleterious blocks (without the beneficial allele) that are formed during the fixation of the beneficial allele. We need to introduce some further notation. Let 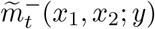 be the mean density of purely deleterious blocks with length at time *t* in the diffusion approximation, starting from a block (*x*_1_, *x*_2_). We will also need to introduce 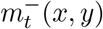, the mean density of blocks of length *y* at time *t* descending from a block of length *x without* the beneficial allele at time 0.

Consider a block (*y*_1_, *y*_2_) that contains the beneficial allele. A crossover occurs to the left of the beneficial allele at rate 2*ρy*_1_, and the resulting block does not contain the beneficial allele with probability 1*/*2. Thus, a purely deleterious block of length *Uy*_1_ is produced at rate *ρy*_1_, with *U* ∼ Uniform(0, 1). Similarly a purely deleterious block of length *Uy*_2_ is produced at rate *ρy*_2_. Once such a block is created, it reproduces according to the rules of the model without a beneficial allele, and with selective effect *σ*_−_ per unit block length.

From this description, the rate at which purely deleterious blocks of length *z* are formed at time *t* − *s* by recombination of a block containing the beneficial allele is

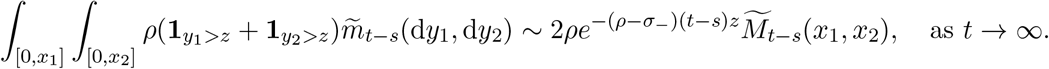

Such a block produces on average 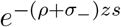 copies of itself between time *t − s* and *t*, and 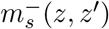 smaller blocks of length *z*′. Therefore, integrating with respect to time from 0 to *t*,

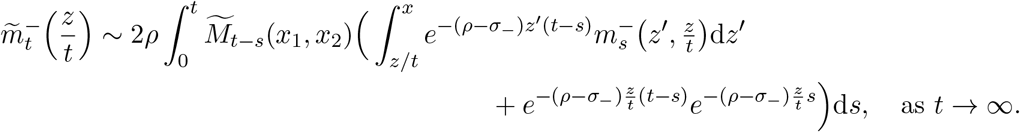

Using that 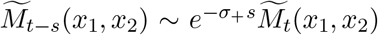 as *t* → ∞ by (A.40) and making a change of variable *y* = *tz*′, the expression above is further approximated by

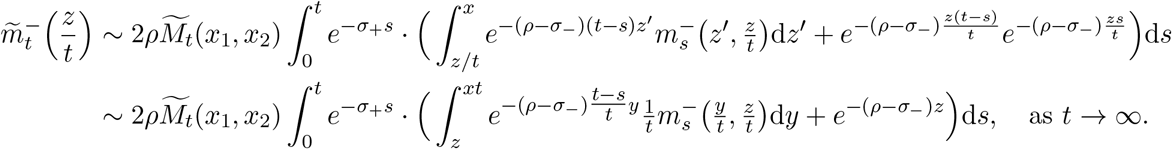

Finally, by (A.30),

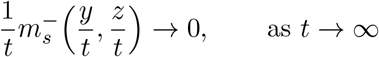

and we deduce that

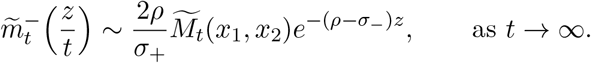

Therefore, the total number of blocks smaller than *z/t* is approximately

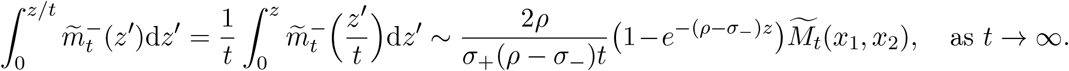

We can re-write this expression in the original units *r* = *ρ/K, s*_+_ = *σ*_+_*/K*, and *s* = *σ /K* at time *t/K* as

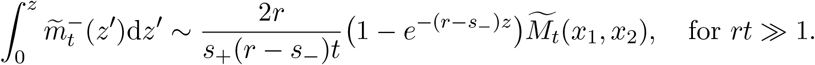

---

**Figure S1:**
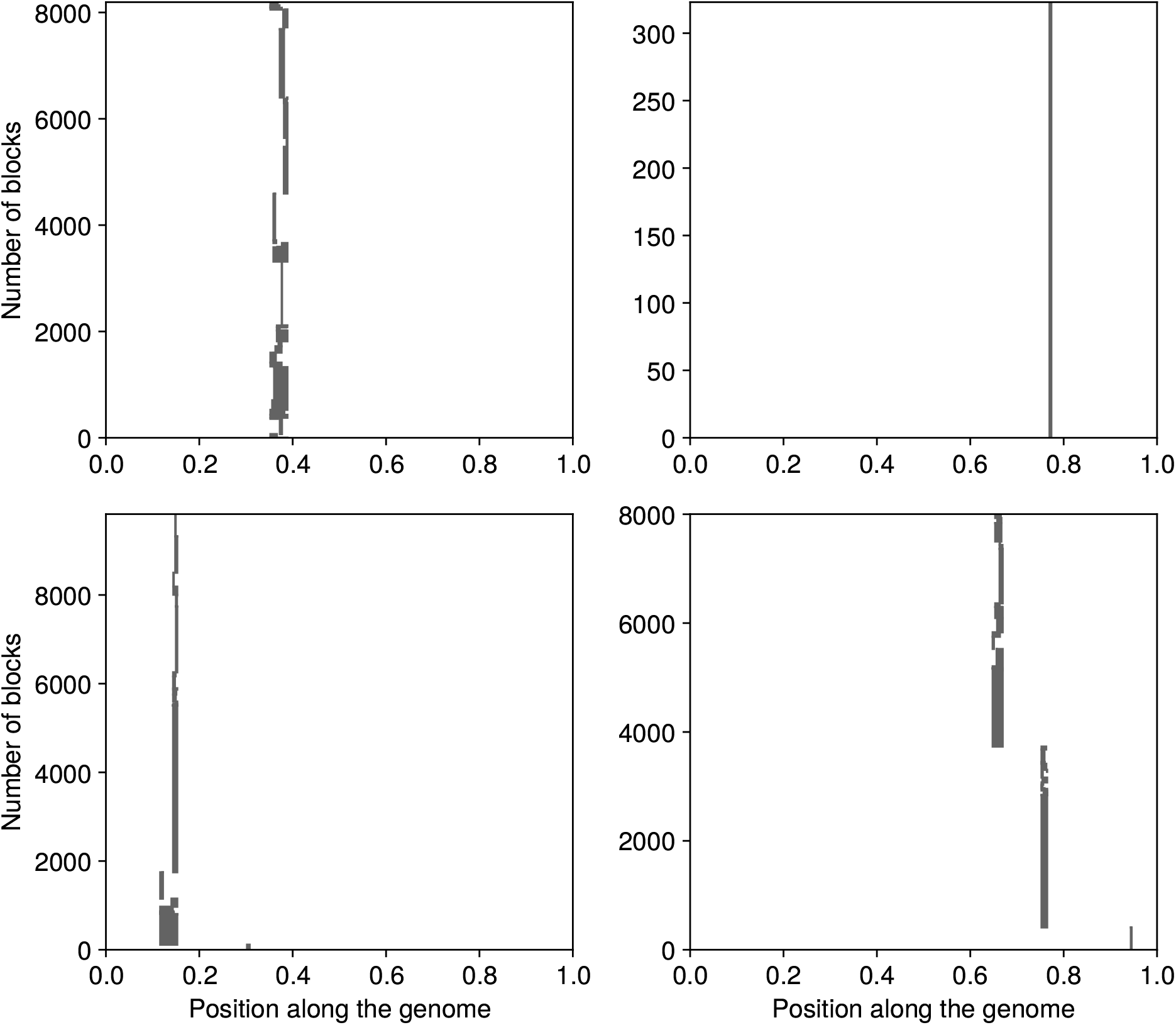
Four realisations of the branching process for *s* = −0.01. All other parameters are described in Figure 1.

**Figure S2:**
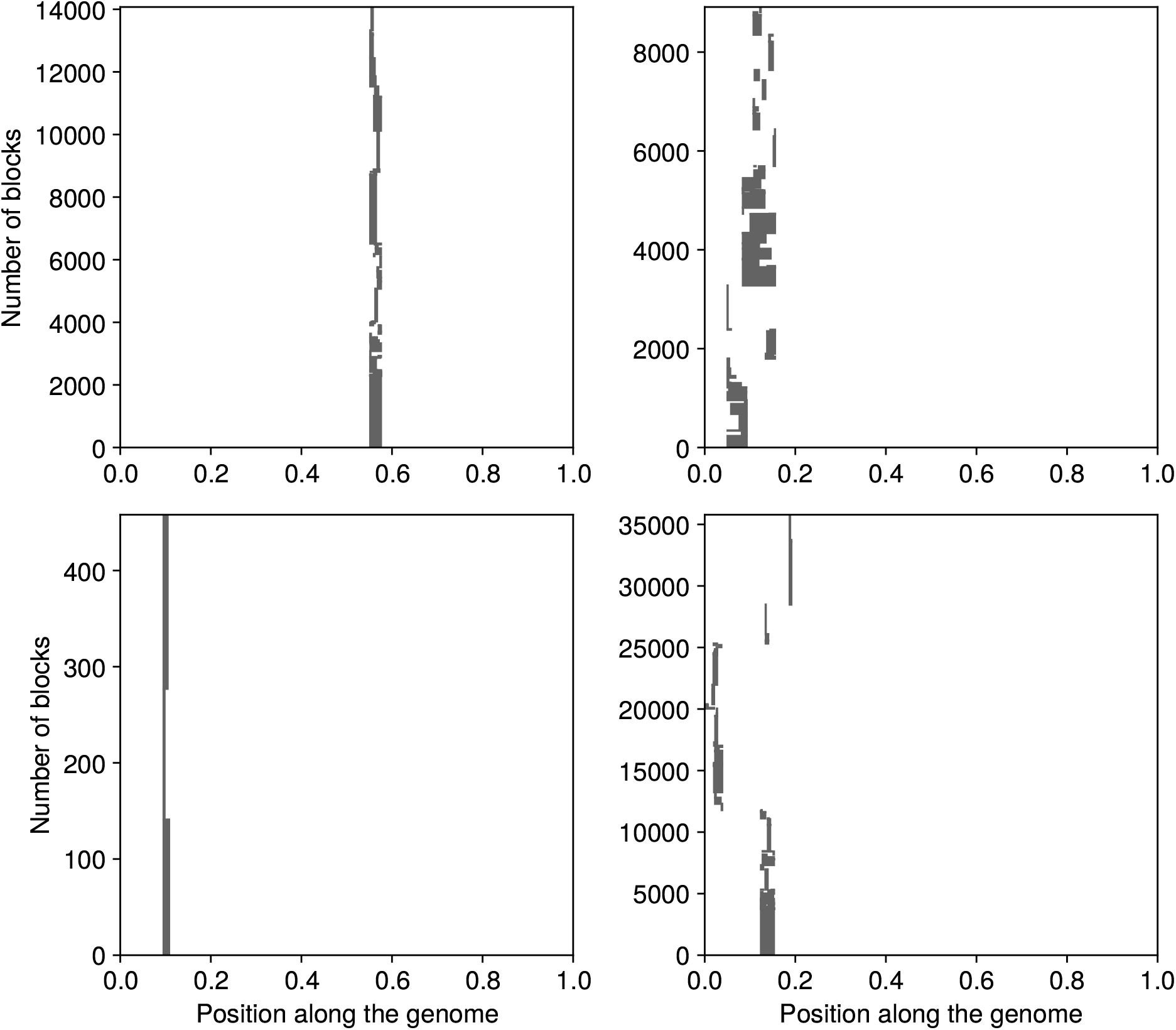
Four realisations of the branching process for *s* = 0. All other parameters are described in Figure 1.

**Figure S3:**
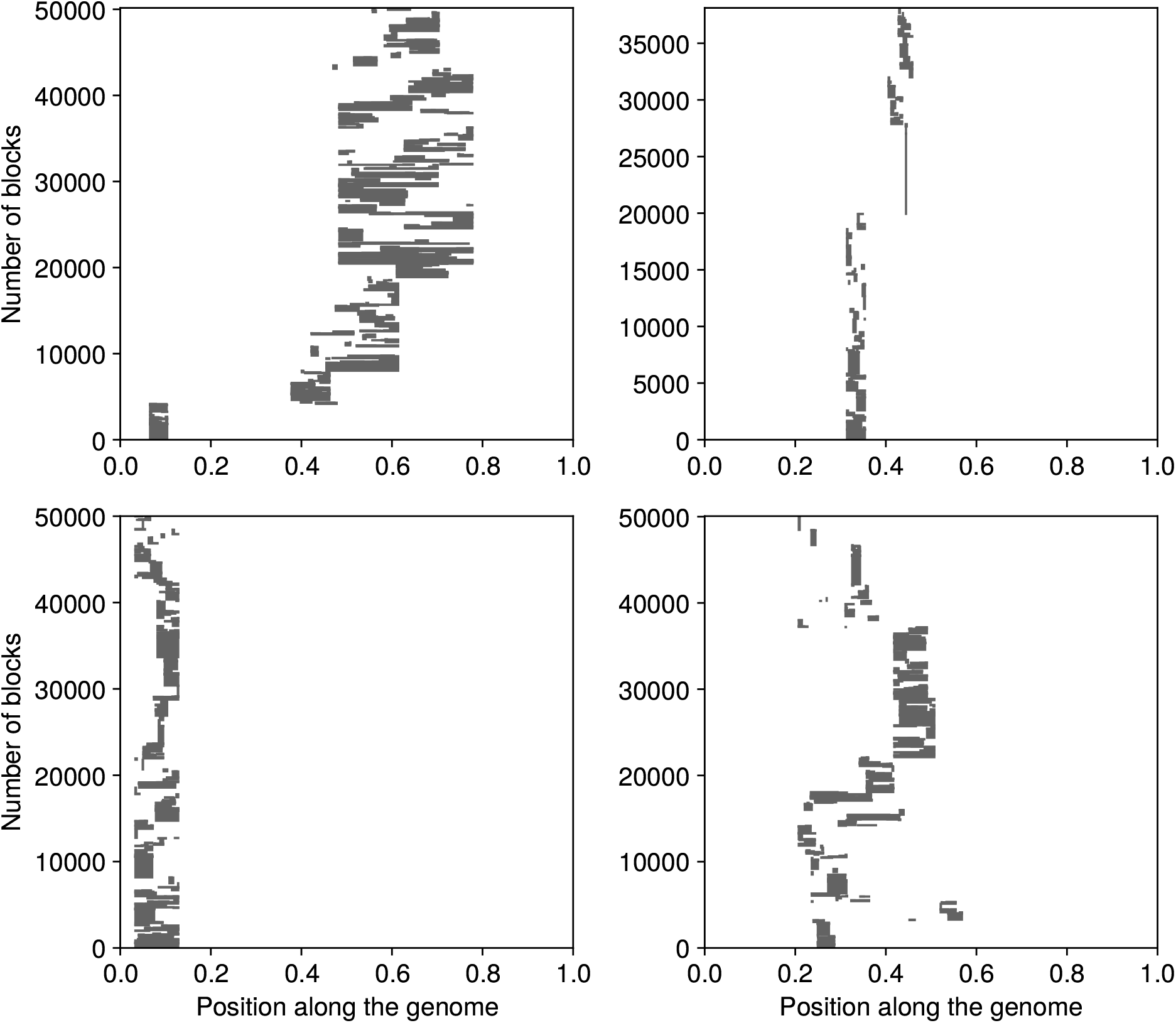
Four realisations of the branching process for *s* = 0.01. All other parameters are described in Figure 1. The process reached 50,000 blocks in three replicates, after 1,971 (top left), 4,085 (bottom left), and 2,701 (bottom right) generations respectively.

**Figure S4:**
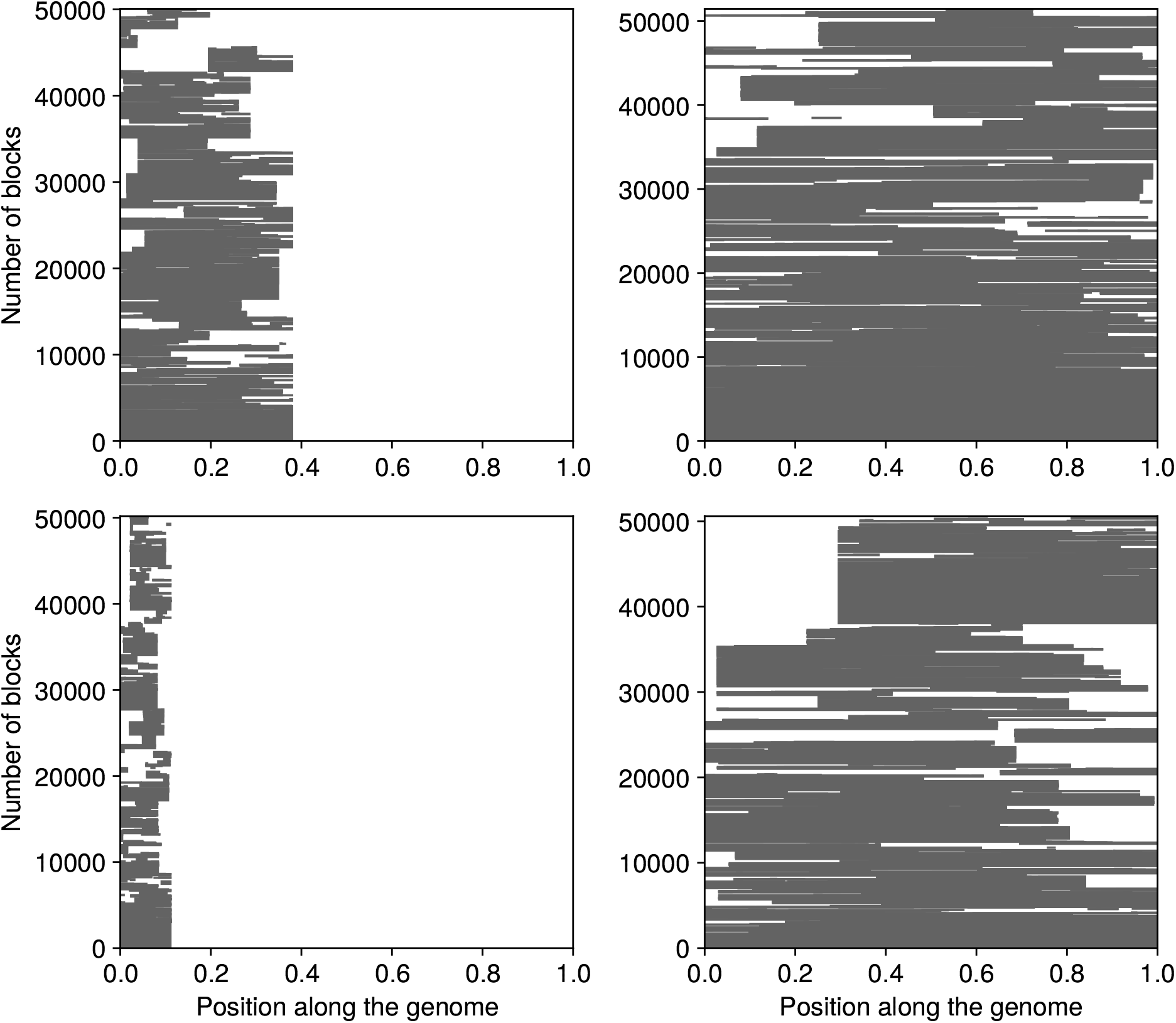
Four realisations of the branching process for *s* = 0.04. All other parameters are described in Figure 1. The process reached 50,000 blocks in all replicates, after 460 (top left), 237 (top right), 1,361 (bottom left), and 264 (bottom right) generations respectively.

**Figure S5:**
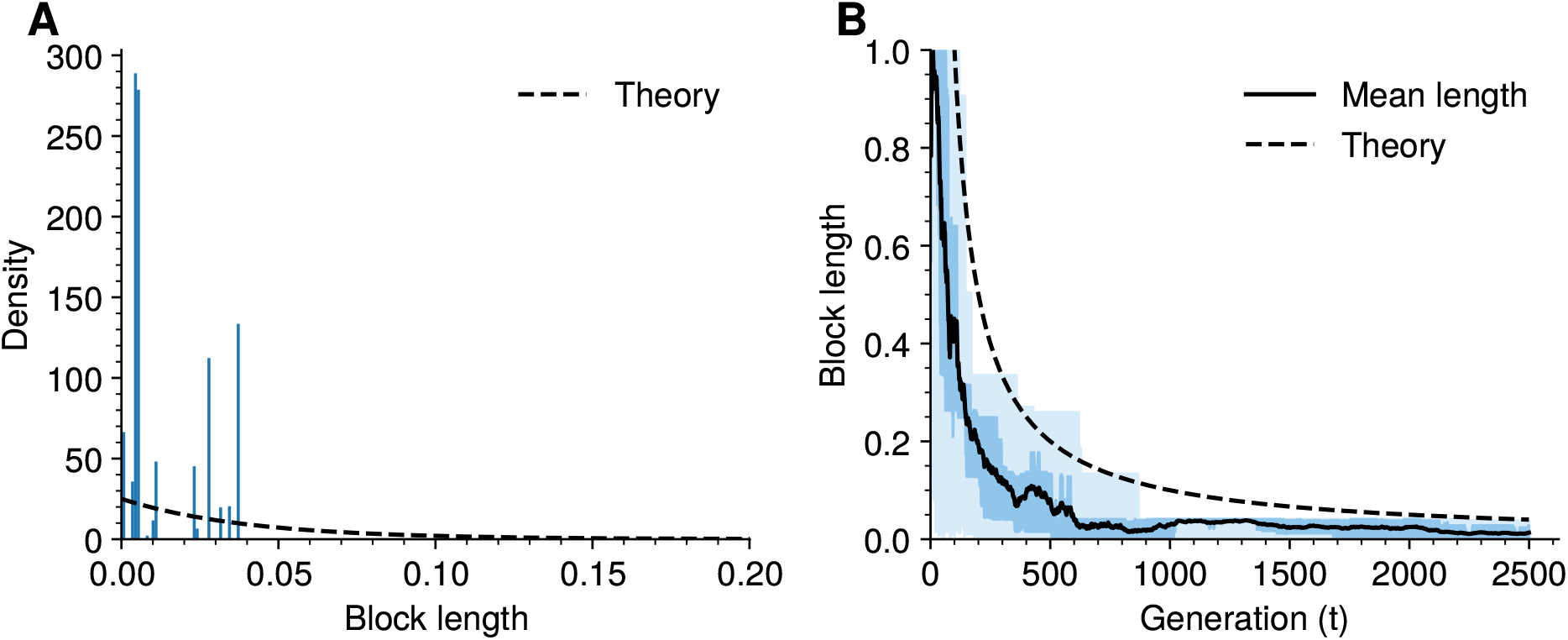
Another independent realisation of the process described in Figure 6. The largest (and average) block length decays fast, probabibly due to an early recombination event. The distribution of block lengths (left) does not converge to a limiting exponential shape but is made of several atoms, corresponding to clones of the same block. This is reminiscent of the example displayed in Figure 1.A for *s <* 0.

## Footnotes

1 As a side note, the solution to (A.11) can be expressed in terms of that to *x* = *x*^*m*^ + *z*, which is the topic of the original article of Lambert (1758). Lambert’s *W* function is a special case that was later studied by Euler (1783).

